# Human blood exposure to *Clostridium perfringens* epsilon toxin may shed light on erythrocyte fragility during active multiple sclerosis

**DOI:** 10.1101/789123

**Authors:** K. Rashid Rumah, Olawale E. Eleso, Vincent A. Fischetti

## Abstract

During active multiple sclerosis (MS), red blood cells (RBCs) harvested from patients reportedly display increased osmotic fragility and increased cellular volume (macrocytosis). The cause of these abnormalities remains unknown. We have previously proposed that *Clostridium perfringens* epsilon toxin (ETX) may be a blood-borne trigger for newly forming MS lesions based on its tropism for blood-brain barrier vasculature and CNS myelin. Recently, Gao et al. have reported that ETX binds to and damages human RBCs, leading to hemolysis. Moreover, the authors suggest that purinergic nucleotide (P2) receptor activation amplifies the hemolytic process. Here, we confirm that ETX indeed causes human-specific RBC lysis. However, our data suggest that the hemolytic process is mediated by metal-catalyzed oxidation of the swell-induced, nucleotide-sensitive ICln chloride channel. We use spectrophotometry, flow cytometry and Western blotting to show that ETX targets human RBCs and T lymphocytes via their shared expression of Myelin and Lymphocyte protein (MAL); a protein shown to be both necessary and sufficient for ETX binding and toxicity. ETX likely triggers T cells to release redox-active heavy metals, Cu^+^ and Fe^3+^, via the lysosomal exocytosis pathway, while RBCs likely release these heavy metals via ETX pore formation within the RBC membrane. Extracellular Cu^+^ and Fe^3+^ may then amplify hemolysis by oxidizing a previously identified heavy metal-binding site within the ICln channel pore, thus deregulating its normal conductance. Elucidating the precise mechanism of ETX-mediated hemolysis may shed light on the underlying etiology of MS, as it would explain why MS RBC abnormalities occur during active disease.

**IMPORTANCE:** During active MS, numerous reports suggest that circulating RBCs are larger than normal and fragment more easily. The exact trigger(s) for these RBC abnormalities and for newly forming MS lesions remains unidentified. We have proposed that ETX, secreted by the gut bacterium *Clostridium perfringens*, may be an environmental trigger for newly forming MS lesions. Indeed, ETX has been shown to breakdown the BBB, enter the brain and damage the myelin sheath. Because ETX is typically spread through the circulatory system, we wished to determine how the toxin affects human blood. Provocatively, there has been a recent report that ETX produces cellular abnormalities in human RBCs, reminiscent of what has been described during active MS. In our study, we sought to elucidate the precise mechanism for how ETX causes RBC damage. In addition to triggering BBB breakdown and CNS demyelination, ETX might also explain why RBCs appear abnormal during MS attacks.

## INTRODUCTION

Although typically considered a disease confined to the central nervous system (CNS), multiple sclerosis (MS) has frequently been shown to cause hematologic abnormalities. To our knowledge, Waksman provided the lone report of T-lymphocyte abnormalities during active MS, which entailed cellular enlargement and a decline in circulating T cell numbers [1]. However, there have been numerous reports of red blood cell (RBC) abnormalities during, and up to one week prior to the onset of neurologic symptoms [2–7]. These decades-old RBC studies have described increased RBC volume (macrocytosis) and increased osmotic fragility, which makes hemolysis more likely to occur. Interestingly, Lewin et al. have recently shed additional light on MS-related RBC abnormalities by showing that free hemoglobin, presumably released after hemolysis, correlates with iron deposition along CNS blood vessels. Iron deposition may lead to neuronal toxicity, axonal loss and the progression from relapsing-remitting MS (RRMS) to secondary-progressive MS (SPMS) [8]. Despite the possible relevance to how MS fundamentally progresses, these hematologic abnormalities remain unexplained.

We have previously suggested that newly forming MS lesions may be triggered by a gut-derived neurotoxin, *Clostridium perfringens* epsilon toxin (ETX), which is disseminated via the circulatory system [9–12]. In support of this theory, Wagley et al. have recently identified serological evidence of prior ETX exposure in MS patients from the United Kingdom [13], complementing what we have observed in an American cohort [9].

*C. perfringens* is an anaerobic, spore-forming, gram-positive bacillus that is sub-classified into seven distinct toxinotypes (A-G) based on differential exotoxin production [14]. *C. perfringens* type A typically colonizes the human gut with a prevalence of 63% among healthy individuals [15], while *C. perfringens* types B and D, the producers of ETX, are commonly found in the intestines of ruminant animals such as sheep, goats, and cattle but rarely in humans [16]. ETX is a potent neurotoxin secreted as a 33 kDa inactive precursor during the logarithmic growth phase of *C. perfringens* in the mammalian intestine. This poorly active precursor is cleaved by gut trypsin, chymotrypsin and several other carboxypeptidases [17]. The ∼27 kDa ETX cleavage product permeablizes the gut epithelium, enters the blood stream and binds to receptors on the luminal surface of brain endothelial cells [9, 16]. Once bound to brain microvessels, ETX oligomerizes and forms a heptameric pore in the endothelial cell plasma membrane. Brain endothelial cell damage leads to breakdown of the BBB [16]. In addition to its known effects on BBB vasculature, ETX has been found to specifically bind to and damage myelin when incubated with mammalian brain slices [11, 18, 19]. This unique ability to interact specifically with the tissues that are damaged in MS, i.e., the BBB and CNS myelin, makes ETX a promising candidate as an environmental MS trigger.

ETX binds to and damages cells that express the tetraspan transmembrane myelin and lymphocyte protein (MAL); relevant cell types containing MAL include the myelin-oligodendrocyte unit, blood-brain barrier endothelial cells and circulating CD4+ T-lymphocytes [10]. Intriguingly, Gao et al. have recently reported that ETX directly binds to the human RBC plasma membrane, matures into the ETX pore-complex and triggers dose-dependent hemolysis, while sparing RBCs from other species. Furthermore, they propose that ETX-mediated hemolysis is amplified by purinergic nucleotide (P2) receptor activation [20].

In this study, we confirm that ETX indeed binds to and damages human RBCs, likely due to their expression of MAL isoform C. However, our data point to a different nucleotide-sensitive system being responsible for amplifying hemolysis. We find that inhibitors of the nucleotide-sensitive, swell-induced chloride channel, ICln, successfully inhibit the hemolytic process.

Because ETX has only recently garnered interest in its potential to cause human disease, there is a paucity of studies published on how it affects human blood. Gaining insight into how ETX triggers human RBC fragility may provide ancillary support for the ETX-MS hypothesis in light of the fact that MS-associated RBC abnormalities remain unexplained. Furthermore, if ETX is ultimately shown to be involved in triggering new MS lesions, understanding the precise mechanism of toxin-triggered hemolysis may open avenues for novel biomarkers capable of predicting the onset of neurologic symptoms, as RBC abnormalities have been shown to precede clinical relapse in some instances. Considering the recent report that hemolysis may play a critical role in the transition from RRMS to SPMS [8], elucidating how hemolysis occurs may allow for clinical interventions aimed at preventing disease progression and permanent neurological decline.

## RESULTS

### Human blood is uniquely sensitive to ETX-mediated hemolysis via MAL expression

We first attempted to repeat the finding by Gao et al. that ETX-mediated hemolysis is specific to human blood [20]. We compared the lytic effect of ETX on human blood vs. blood from ruminant animals such as sheep, goats and cows, as these animals are the natural hosts for ETX-secreting *C. perfringens* strains [16]. We also tested guinea pig blood as a non-human, non-ruminant control. Highly purified epsilon protoxin (protoETX), as determined by SDS-PAGE (Supplemental Fig 1) was trypsin activated, and time-course analysis showed that this activated ETX (15nM) caused hemolysis in PBS-washed whole human blood. However, blood harvested from non-human species was completely refractory (Fig 1a). Importantly, we also determined that trypsin activation was necessary for hemolytic activity, as non-activated protoETX failed to trigger hemolysis (Supplemental Fig 2). Moreover, inhibition of activated ETX with an anti-ETX neutralizing antibody, JL008, inhibited hemolysis in a dose-dependent manner (Supplemental Fig 3). We also confirmed that ETX directly binds human RBCs by incubating cells harvested from human, rhesus macaque, sheep and rat with non-activated protoETX (50nM), which is capable of cellular binding, but is incapable of forming pores within the cell membrane [16]. We visualized human-specific ETX binding by flow cytometry using custom anti-ETX antibody, JL001.2 (Fig 1b).

**Fig 1.**
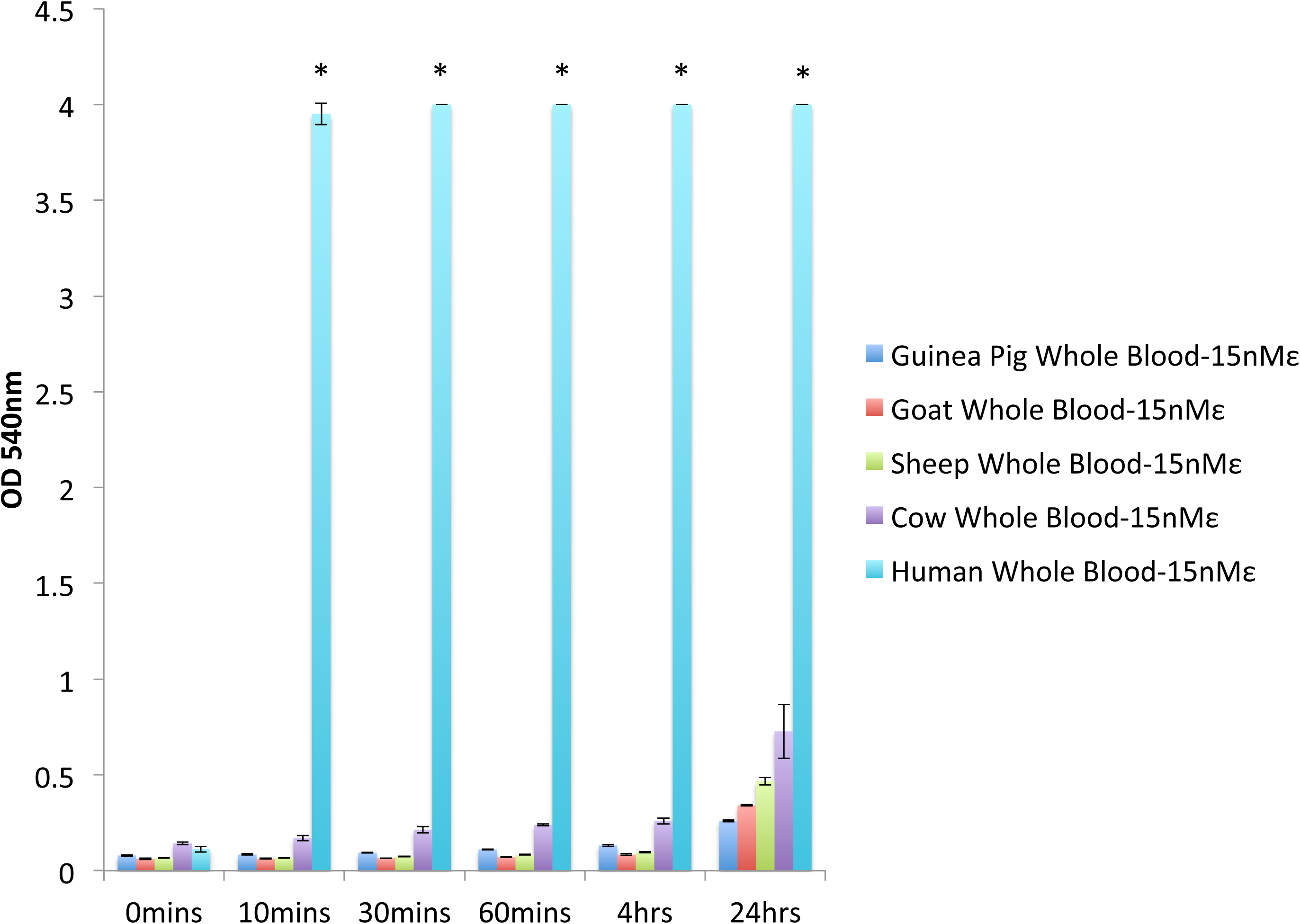
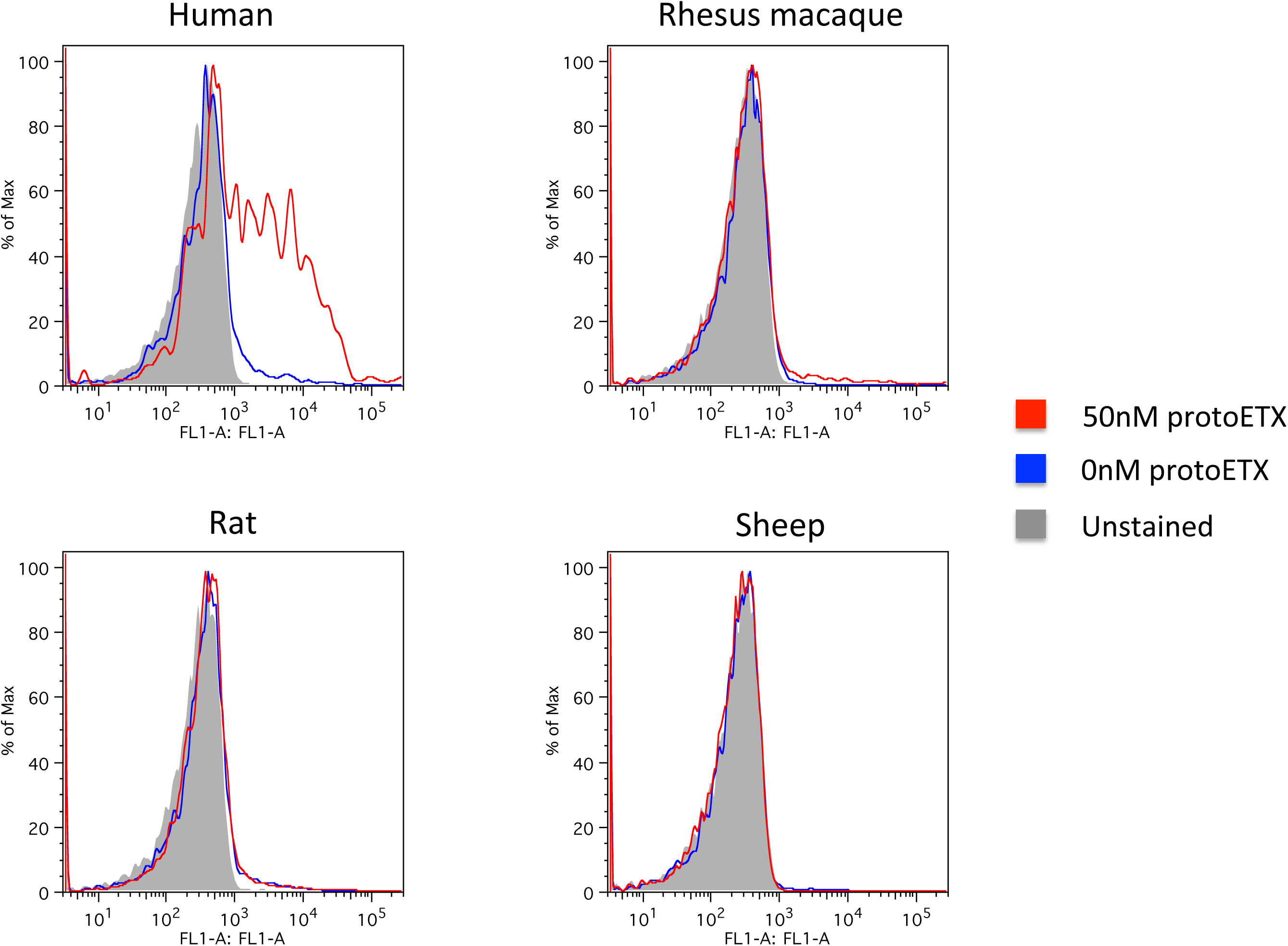
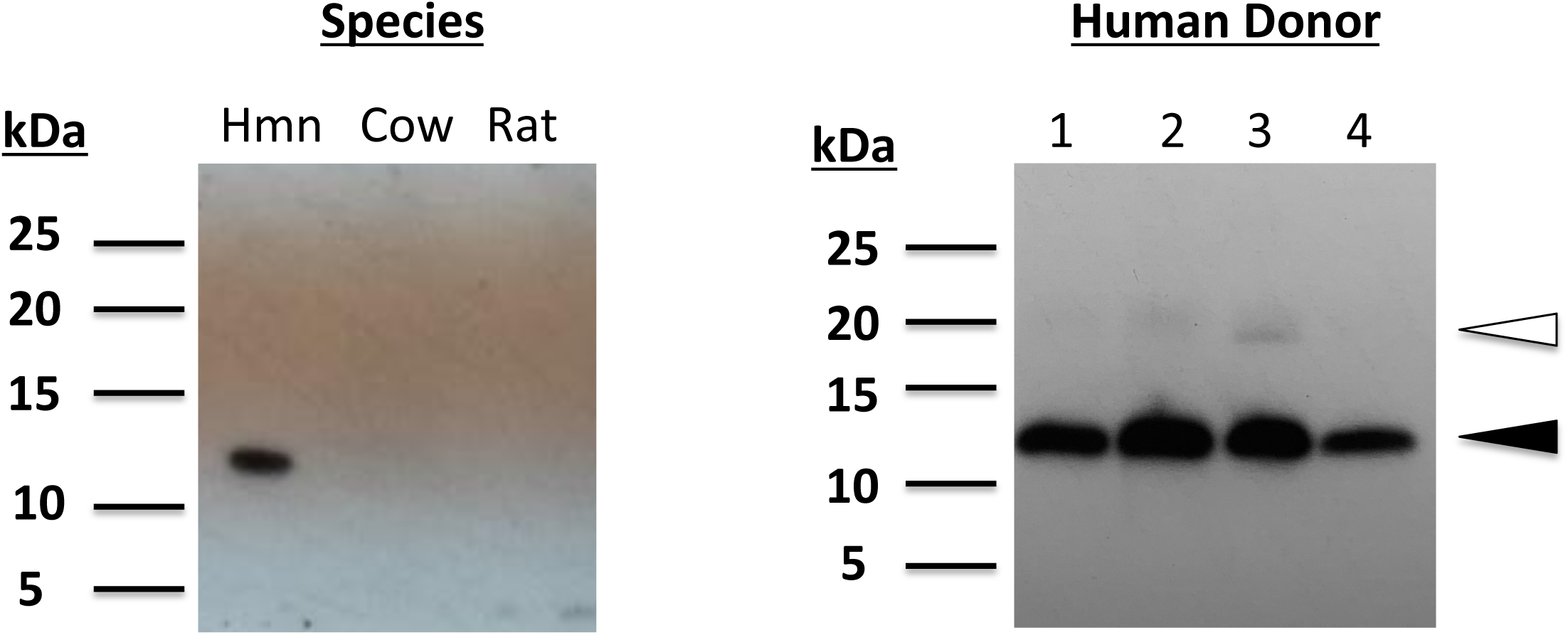
ETX causes human-specific hemolysis via human RBC MAL expression. **a)** Washed whole human, cow, sheep, goat and guinea pig blood was suspended in PBS and incubated with 15nM ETX at 37°C, and sampled over 24 hours. Data shown are from experiments performed in triplicate. Error bars represent standard deviations, and asterisks indicate that results are statistically significant compared with refractory guinea pig blood (dark blue); Student’s *t*-test, \**P* < 0.0001. **b)** Human, rhesus macaque, rat and sheep RBCs were incubated with non-toxic protoETX (50nM) for 2 hours at 4°C. Toxin binding was detected via flow cytometry using custom anti-ETX antibody, JL001.2. Data shown are from a single experiment and are representative of 3 independent experiments, using 3 distinct blood donors. **c)** Human, cow and rat RBC membranes were analyzed for MAL expression by Western blot (left). RBC membranes harvested from additional human donors were also analyzed (right). The open arrowhead signifies full length MAL, while the closed arrowhead signifies the shortened MAL isoform.

Our previous work has shown that lipid raft protein, MAL (17 kDa, predicted), is both necessary and sufficient for ETX binding and toxicity, thus we sought to determine if human RBCs express MAL. Consistent with MAL’s proposed role in ETX toxicity and with previously published RBC membrane proteomic analysis [21], we corroborated that human RBCs indeed express MAL; specifically a shortened isoform predicted to be 11kDa, MAL isoform C (Fig 1c). RBCs from refractory species, cow and rat, were both negative for MAL expression. However, we cannot exclude the possibility that the recognition antibody used may be specific for human MAL. Interestingly, there seemed to be a trace expression of full length MAL (open arrowhead) in addition to the dominant band appearing for shortened MAL isoform C (closed arrowhead).

### P2 receptor agonists and antagonists both inhibit ETX-mediated hemolysis

While assessing the possible involvement of the P2 purinergic nucleotide receptors in ETX-mediated hemolysis (as suggested by Gao et al.), we first exposed PBS-washed whole human blood to: i) the irreversible P2 receptor antagonist, oxidized ATP (oxATP); ii) P2 receptor agonists ATP and GTP and; iii) UMP, as a poorly interacting nucleotide control, before treatment with ETX [22]. Surprisingly, we found that both P2 agonists and P2 antagonists inhibited ETX-mediated hemolysis (Fig 2a). Furthermore, we found that overnight incubation with oxATP and subsequent washout of this irreversible P2 inhibitor prior to ETX treatment largely abolished its anti-hemolytic effect (Fig 2b). These results led us to search for alternative pathways capable of amplifying ETX-mediated hemolysis in a nucleotide-sensitive fashion.

**Fig 2.**
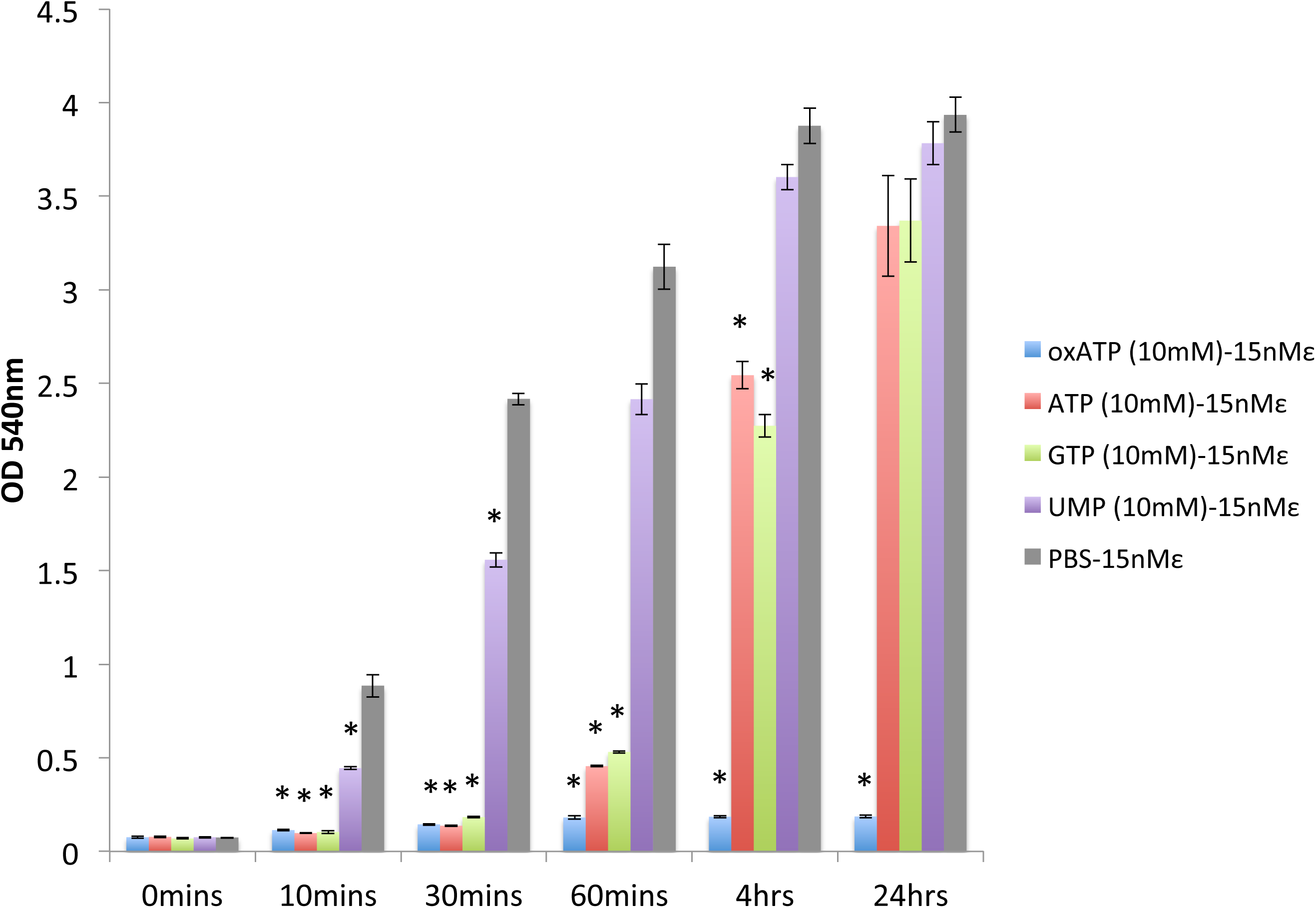
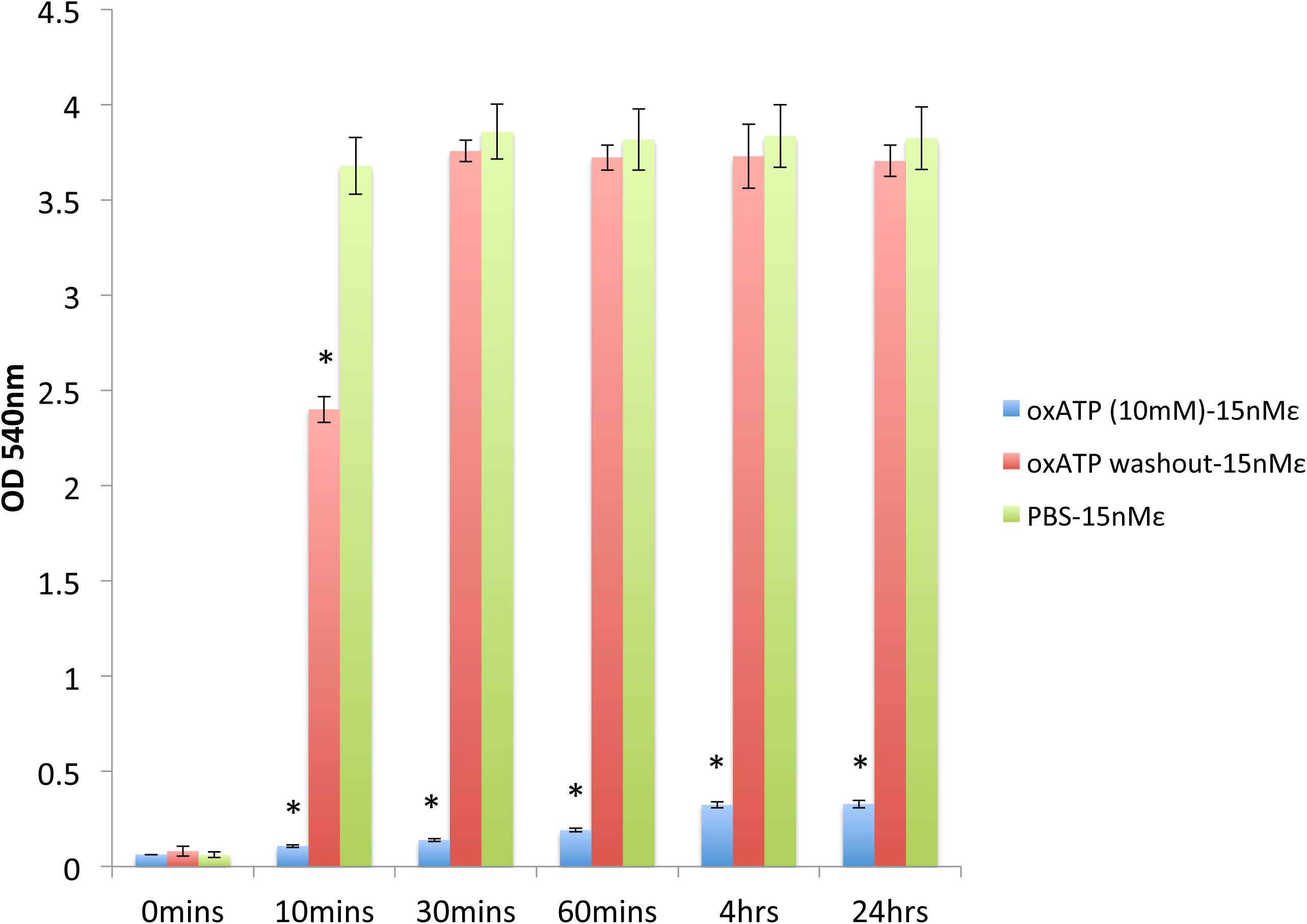
Extracellular nucleotides inhibit ETX-mediated hemolysis. **a)** Washed whole human blood was pre-incubated with oxidized ATP (oxATP), ATP, GTP and UMP (10mM each) prior to ETX exposure (15nM) at 37°C, and sampled over 24 hours. Data shown are from experiments performed in triplicate. Error bars represent standard deviations, and asterisks indicate that results are statistically significant compared with PBS vehicle control (gray); Student’s *t*-test, \**P* < 0.0002. **b)** Washed whole human blood was exposed to the irreversible P2 receptor inhibitor, oxATP, overnight at 37°C. OxATP was either allowed to remain in the suspension buffer (oxATP 10mM) or washed out. All samples were then incubated with ETX (15nM) at 37°C, and sampled over 24 hours. Data shown are from experiments performed in triplicate. Error bars represent standard deviations, and asterisks indicate that results are statistically significant compared with PBS vehicle control (green); Student’s *t*-test, \**P* < 0.0002.

### O_2_ and redox-active heavy metals, Cu^+^ and Fe^3+^, are involved in ETX-mediated hemolysis

A serendipitous finding, observed upon exposing ETX-treated blood to different experimental conditions, was that atmospheric oxygen is required for hemolysis. One concern was that anaerobiosis may be blocking a pathway crucial to ETX-mediated hemolysis that is dependent on the electron-transport chain and oxidative phosphorylation. To address this, we compared the hemolytic effect of ETX under aerobic, anaerobic and sodium azide (10mM) conditions. Sodium azide was used as an alternative method to inhibit oxidative phosphorylation in lieu of atmospheric oxygen depletion. The results revealed that ETX-mediated hemolysis was inhibited under anaerobic conditions, but not by oxidative phosphorylation inhibition via sodium azide blockade (Fig 3a).

**Fig 3.**
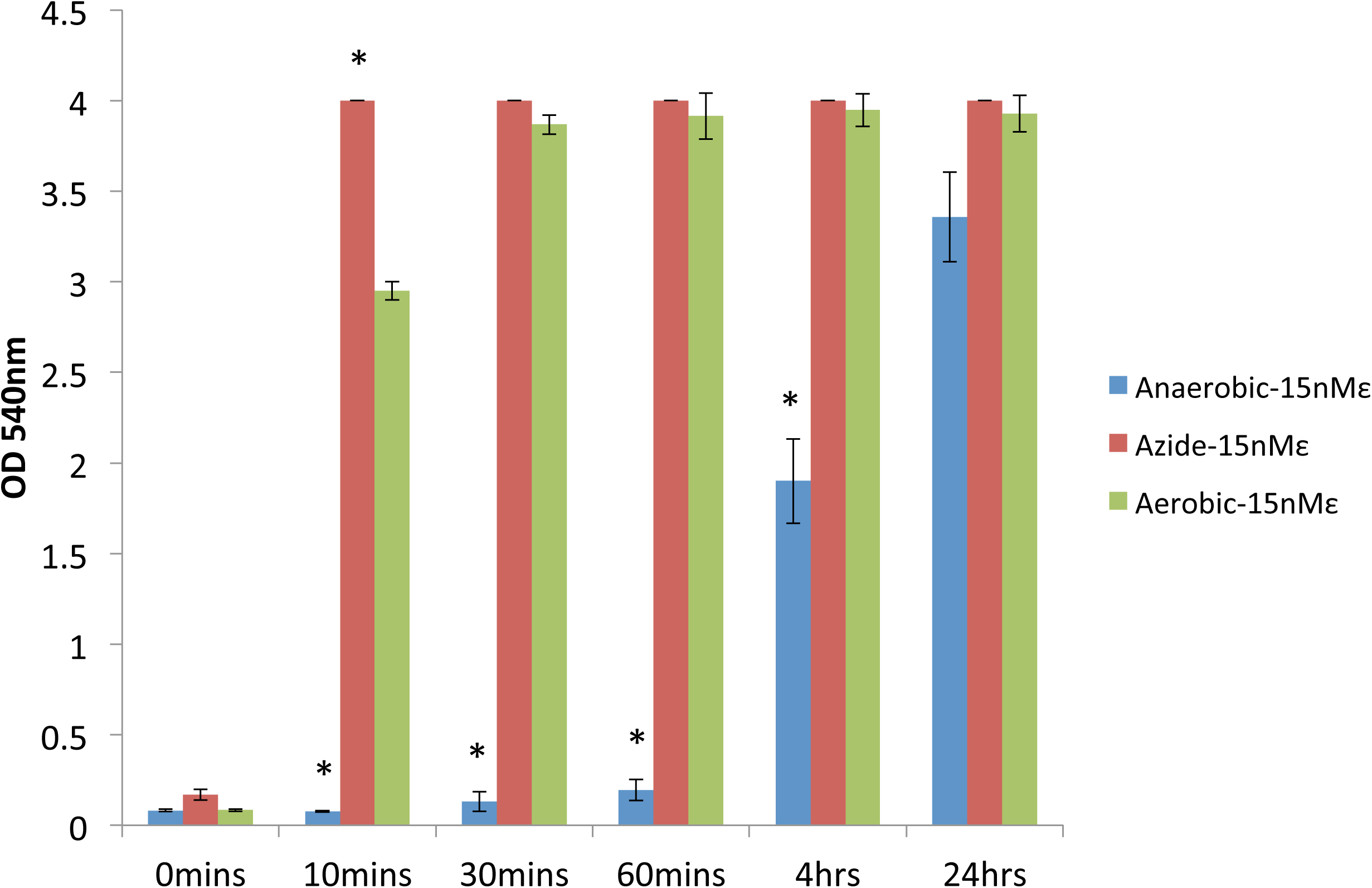
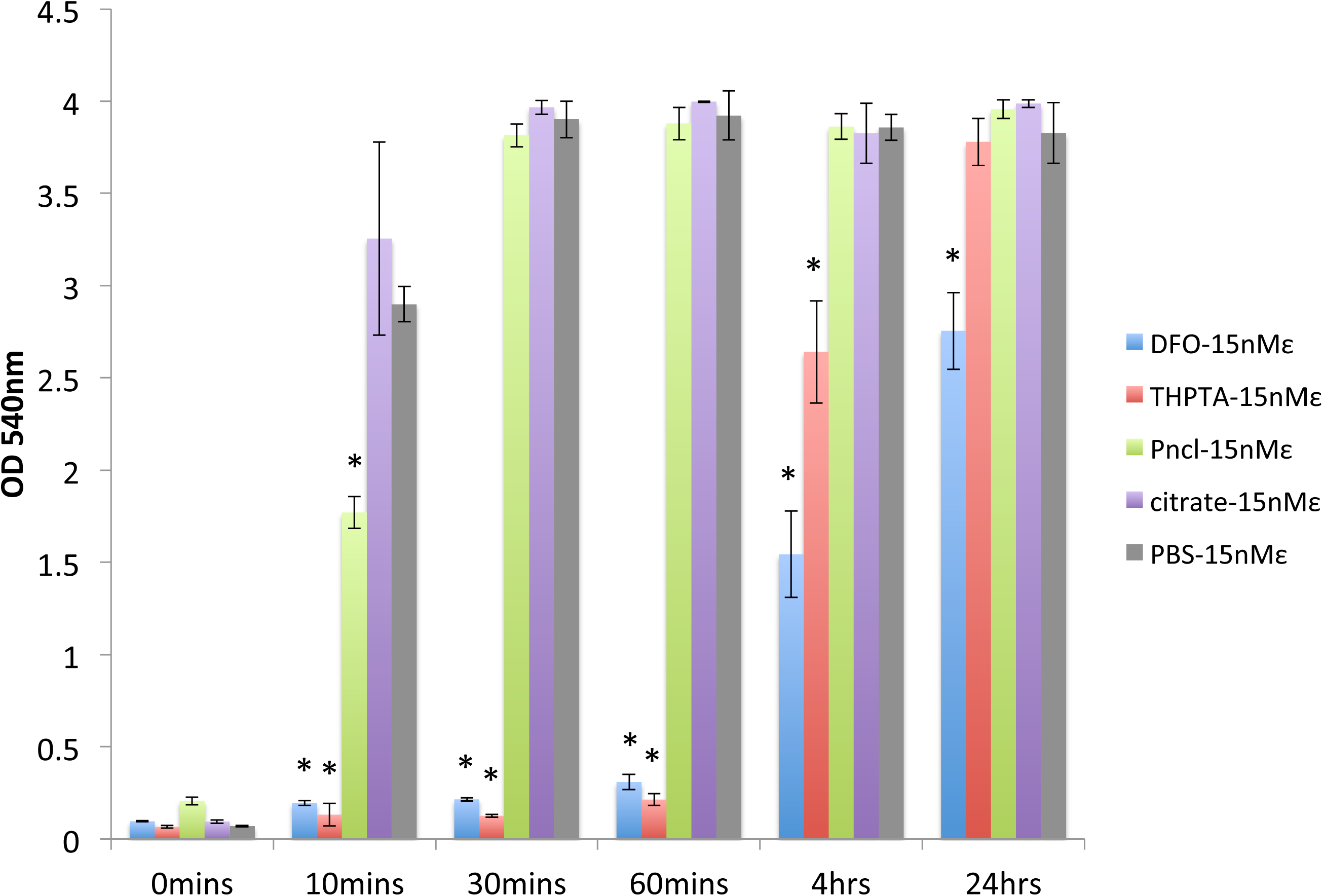
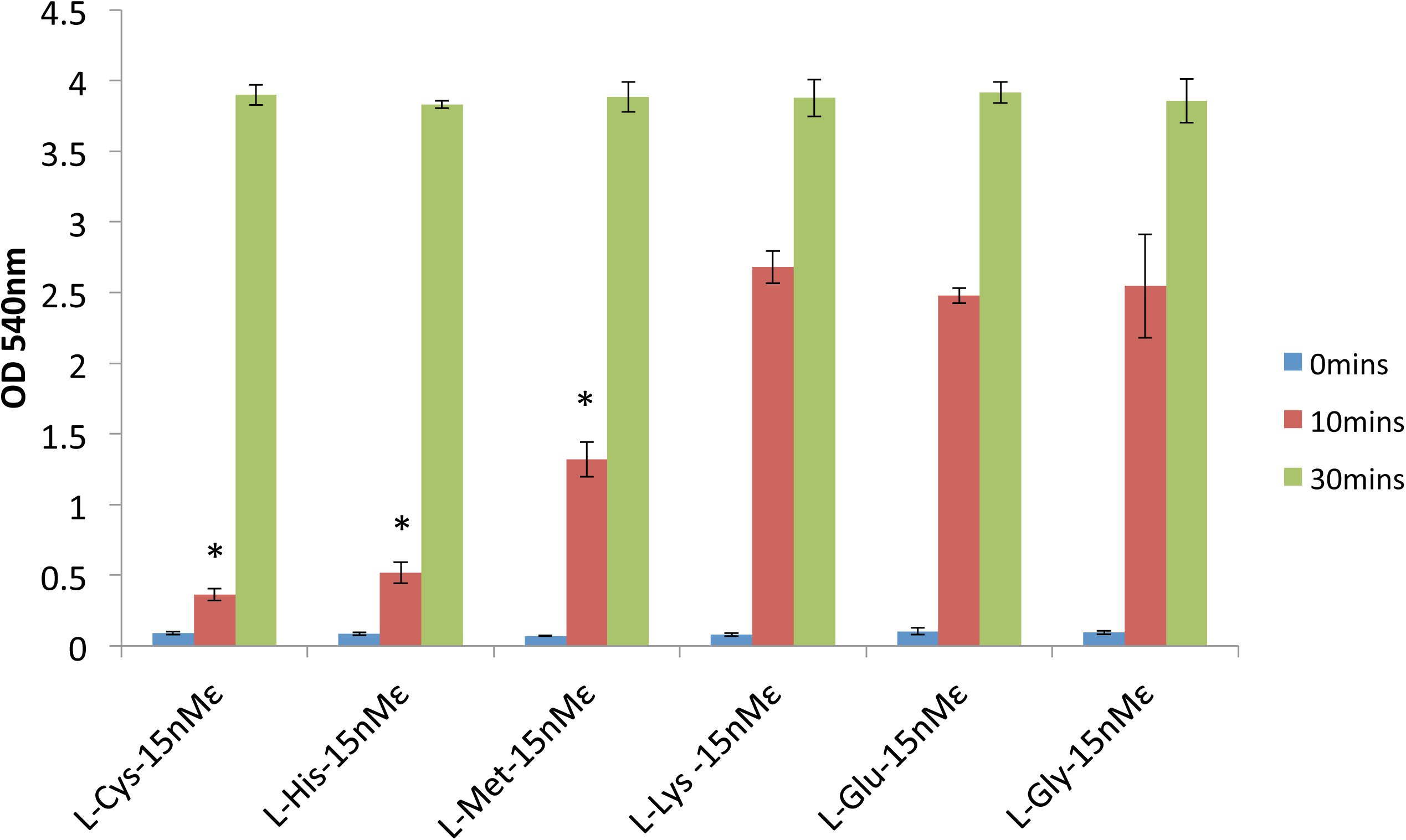
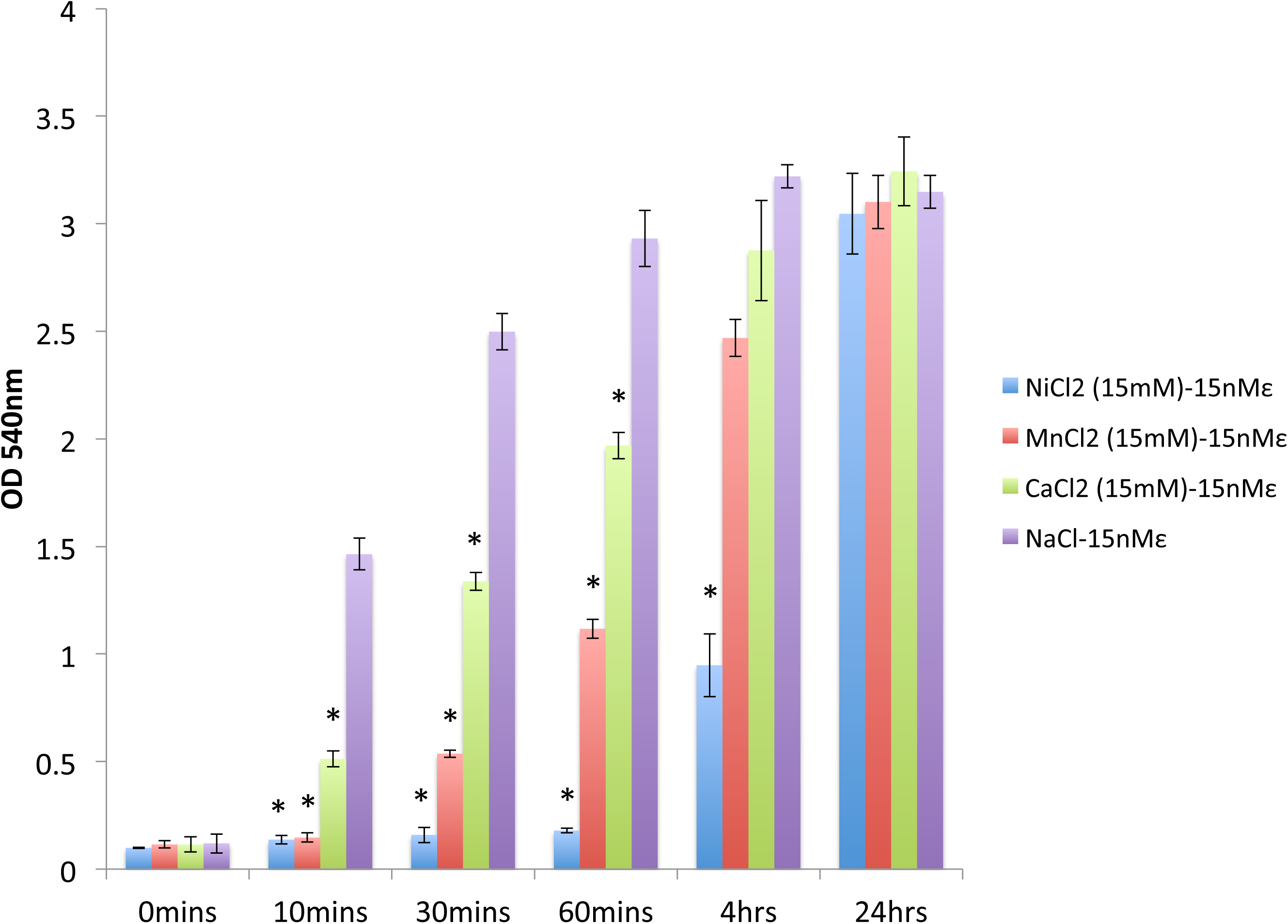
Oxygen and redox-active transition metals, Cu^+^ and Fe^3+^, are integral to ETX-mediated hemolysis. **a)** Washed whole human blood was suspended in PBS and exposed to 15nM ETX at 37°C under anaerobic conditions, aerobic conditions or co-incubated with sodium azide (10mM) and sampled over 24 hours. Data shown are from experiments performed in triplicate. Error bars represent standard deviations, and asterisks indicate that results are statistically significant compared with aerobic control (green); Student’s *t*-test, \**P* < 0.0001. **b)** Washed whole human blood was exposed to ETX (15nM) at 37°C over 24 hours, and co-incubated with metal chelators (5mg/mL each), deferoxamine (DFO), THPTA, penicillamine (Pncl), citrate or PBS buffer control. Data shown are from experiments performed in triplicate. Error bars represent standard deviations, and asterisks indicate that results are statistically significant compared with PBS vehicle control (gray); Student’s *t*-test, \**P* < 0.0003. **c)** The inhibitory effects of endogenous metal chelating amino acids (5mg/mL each), cysteine, histidine and methionine were compared to that of non-chelating amino acids (5mg/mL each), lysine, glutamate and glycine. All samples were exposed to ETX (15nM) at 37°C and sampled over 30 mins. Data shown are from experiments performed in triplicate. Error bars represent standard deviations, and asterisks indicate that results are statistically significant compared with glycine (farthest right); Student’s *t*-test, \**P* < 0.01. **d)** Washed whole human blood was suspended in isotonic saline with or without divalent cations (15mM each), Ni^2+^ or Mn^2+^ (redox-silent transition metals) or Ca^2+^ (alkaline earth metal), and assessed for inhibition of ETX-mediated hemolysis (15nM ETX at 37°C) over 24 hours. Data shown are from experiments performed in triplicate. Error bars represent standard deviations, and asterisks indicate that results are statistically significant compared with NaCl vehicle control (green); Student’s *t*-test, \**P* ≤ 0.0003.

Considering the involvement of ambient O_2_, we explored various sources of cellular free-radial generation. A potent way for macromolecules to be specifically targeted and damaged in an oxygen-dependent manner is by metal-catalyzed oxidation, where redox-active transition metals coordinate with metal-binding amino acid residues, e.g., cysteine, histidine and methionine, and locally generate reactive oxygen species (ROS) [23]. Along these lines, we pre-incubated ETX-treated human blood with the hydrophilic Cu^+^ chelator, tris-hydroxypropyltriazolylmethylamine (THPTA) and the hydrophilic Fe^3+^ chelator, deferoxamine (DFO), both of which significantly inhibited ETX-mediated hemolysis. In contrast, the Cu^2+^ chelator, penicillamine (Pncl), and the non-specific metal chelator, citrate, were not effective at inhibiting hemolysis (Fig 3b). As an alternative to “artificial” metal chelators, we also exposed ETX-treated blood to “endogenous” metal chelators, such as the heavy metal-coordinating amino acids, cysteine, histidine and methionine [24], each of which transiently inhibited ETX-mediated hemolysis, cysteine > histidine > methionine. However, the control amino acids, lysine, glutamate and glycine were significantly less effective (Fig 3c).

A metal-coordinating protein involved in how the RBC membrane interacts with liberated Cu^+^ and Fe^3+^ is likely to bind more than one metallic species, and with varying affinities. This notion led us to hypothesize that pretreatment of human blood with redox-silent transition metals, such as Ni^2+^ and Mn^2+^, might protect RBCs from ETX-mediated hemolysis by competing for crucial metal-binding sites on the RBC membrane. Indeed, we observed that both Ni^2+^ and Mn^2+^ significantly inhibited ETX-mediated hemolysis, Ni^2+^ > Mn^2+^. Ca^2+^ also exhibited inhibitory properties, but to a lesser extent when compared to the redox-silent transition metals (Fig 3d).

### The heavy metal binding, nucleotide-sensitive ICln chloride channel amplifies ETX-mediated hemolysis

The sensitivity of ETX-mediated hemolysis to the presence of nucleotides, ambient oxygen and the extracellular chelation of redox-active transition metals led us to search the literature for a surface molecule, which may be sensitive to these stimuli. We identified the ubiquitously expressed, outwardly-rectifying ICln chloride channel as a lead candidate. Under normal conditions, the ICln protein is found to be closely associated with the actin cytoskeleton abutting the inner leaflet of the plasma membrane [25]. Upon cellular swelling, as would be expected from ETX pore formation, ICln rapidly inserts into the plasma membrane forming a channel that causes intracellular chloride to flow out of the cell, against its diffusion gradient. This outward rectification allows the cell to decrease its volume, thus avoiding osmolysis [26]. Remarkably, a key histidine residue, His64, resides within the ICln pore. His64 coordinates with Ni^2+^ to alter ICln’s normal conductance [27]. We hypothesize that this regulatory, heavy metal-binding residue may be the site at which redox-active Cu^+^ and Fe^3+^, previously liberated by initial ETX pore formation, bind to and damage this volume-regulating channel. Of note, ICln also possesses distinct Ca^2+^ binding sites, located near the extracellular opening of the channel that allow for conductance inhibition by extracellular Ca^2+^ [27].

An alternate name for ICln is chloride channel nucleotide sensitive 1A (CLNS1A). As the name suggests, the presence of extracellular nucleotides also regulates ICln conductance, i.e., ATP and GTP inhibit ICln [28]. To further investigate ICln’s potential role in ETX-mediated hemolysis, we tested other distinct classes of ICln inhibitors, namely the chromones (cromolyn and nedocromil) [29], and the cyclamate anion [30]; each of which successfully inhibited ETX-mediated hemolysis (Figs 4a and 4b). We tested each class of ICln inhibitor in an aggregate experiment to provide a comparative analysis (Fig 4c). Please note that we omitted the anti-retroviral nucleoside analogue ICln inhibitors, acyclovir and AZT [31] due to poor solubility relative to the other ICln inhibitors. Interestingly, when we assessed RBC ICln expression via Western blot, we observed the occasional appearance of an ICln doublet (open arrowhead), consistent with what has been observed in previous studies [25]. However, the doublet did not correlate with changes in the ETX concentration (Fig 4d).

**Fig 4.**
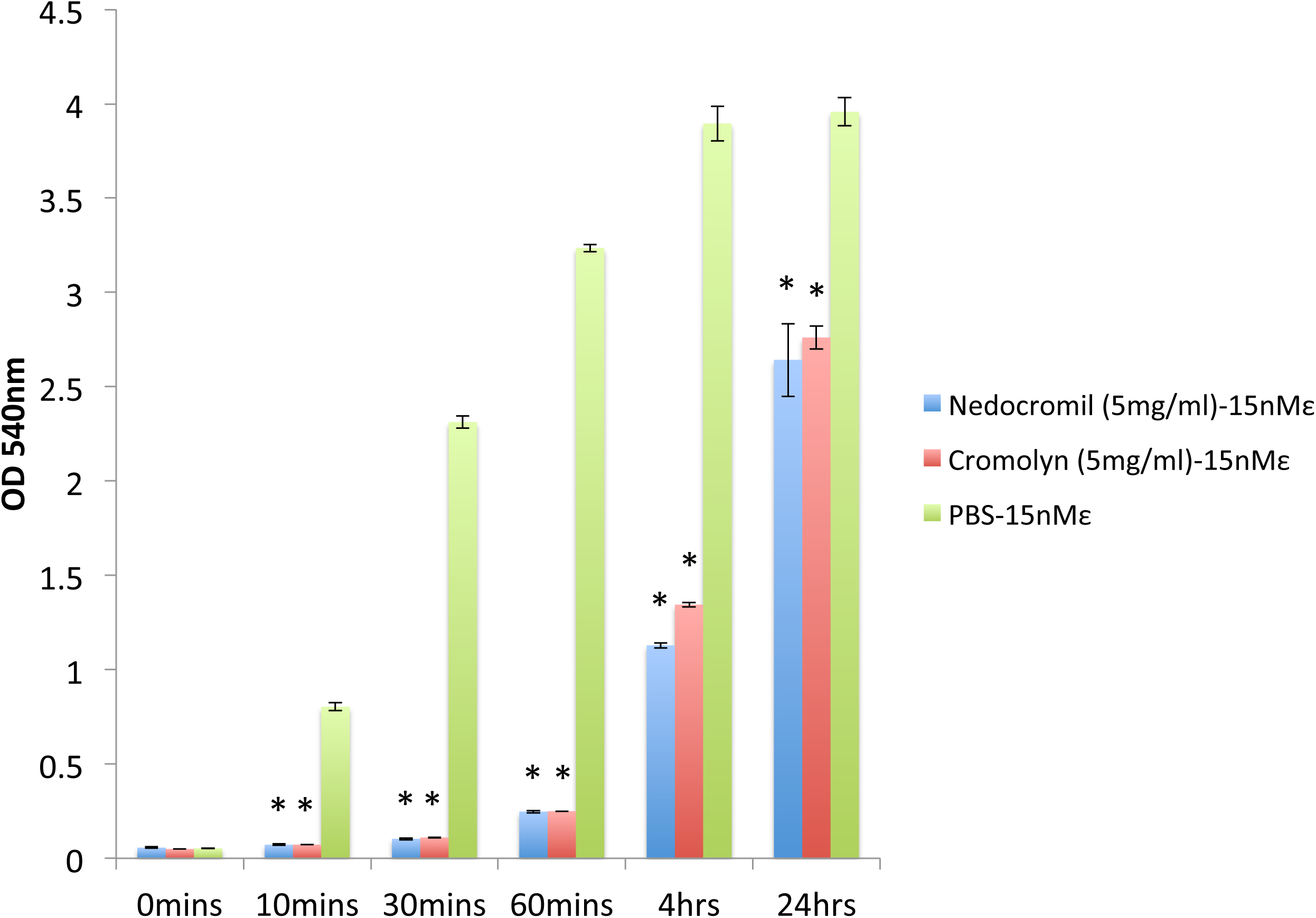
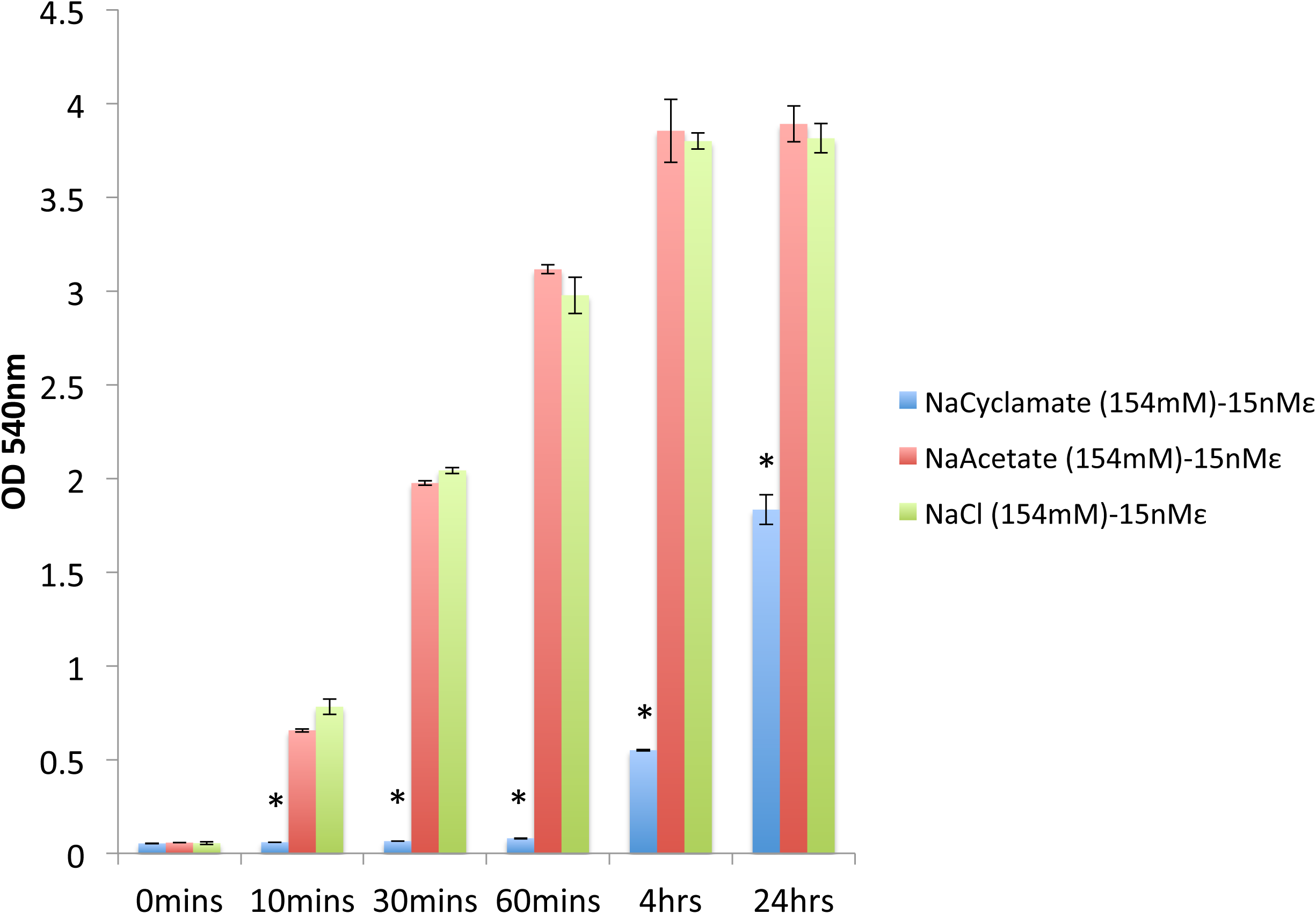
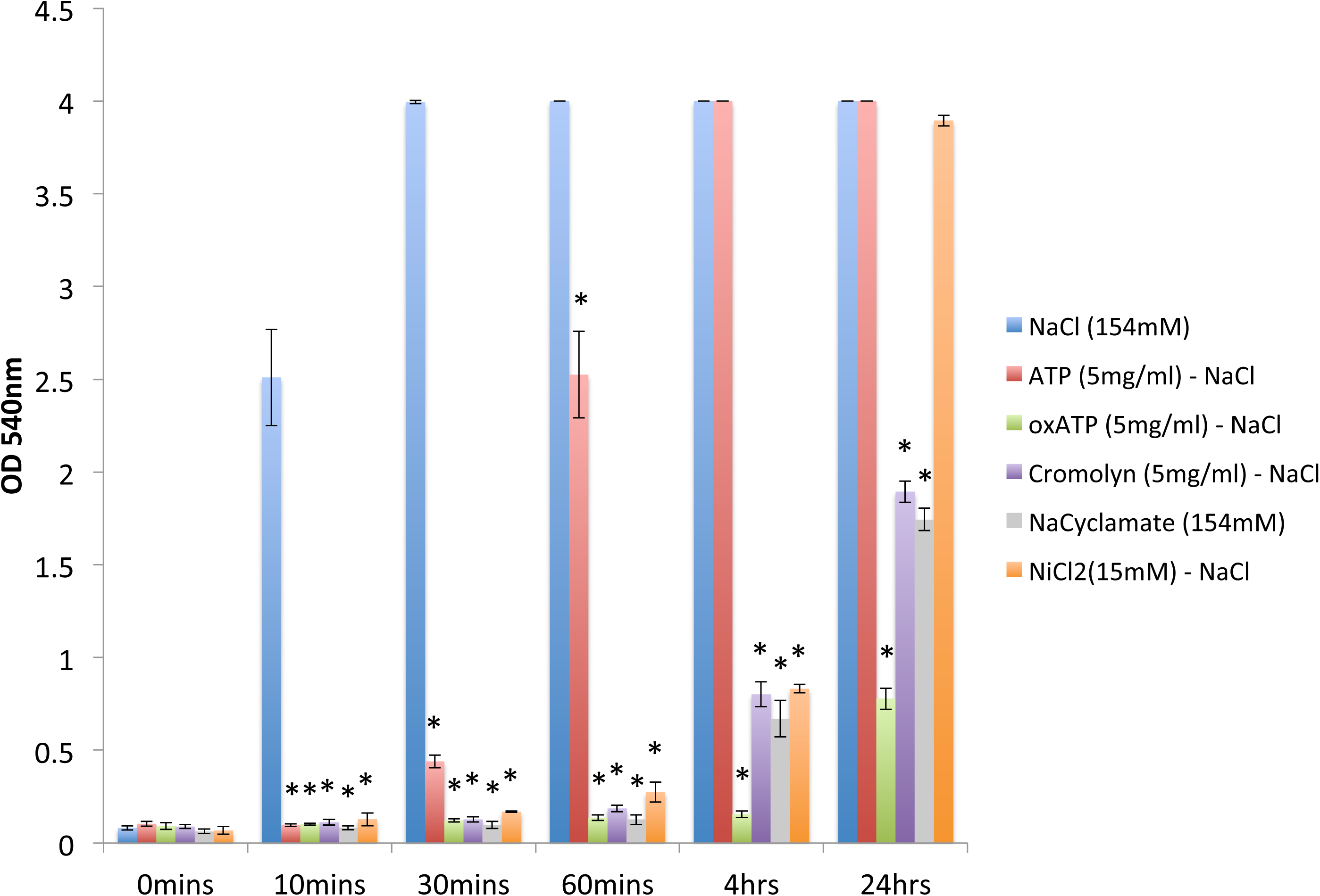
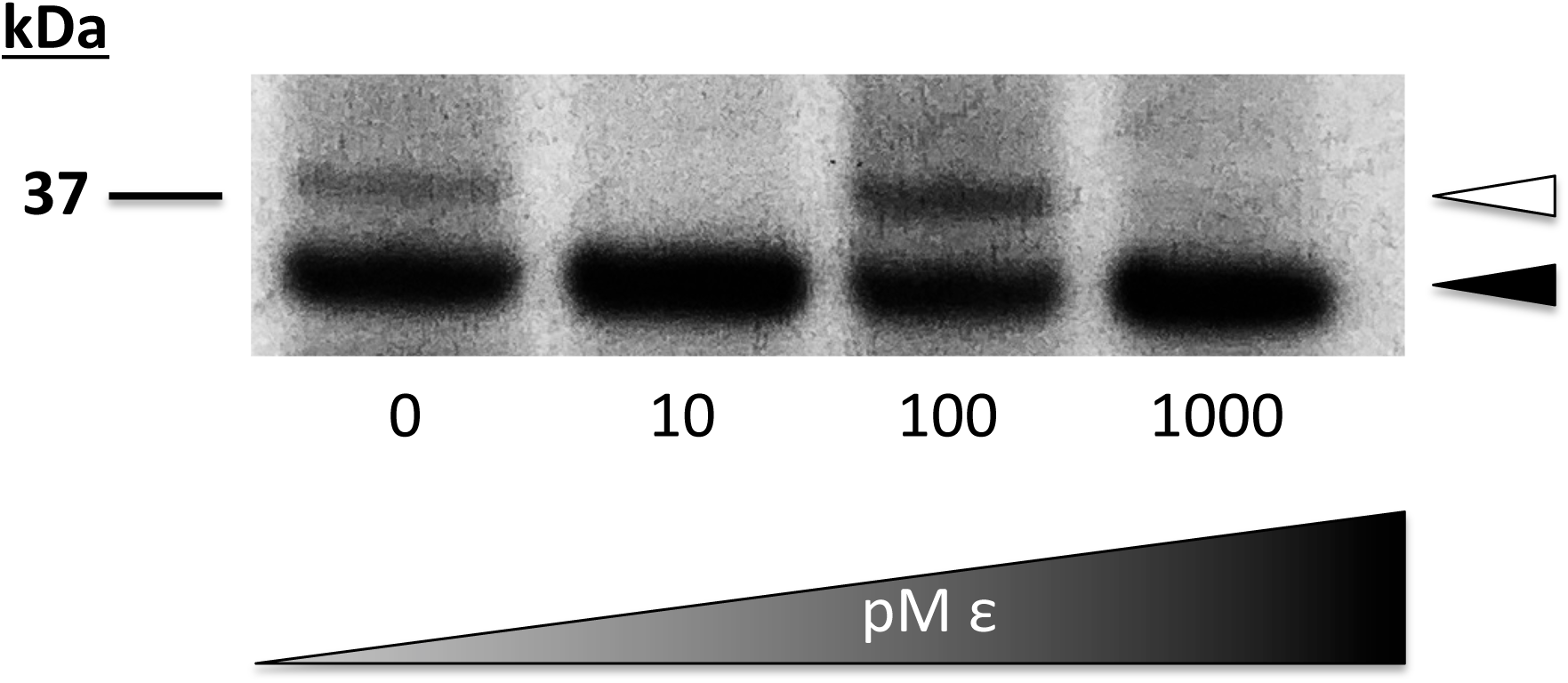
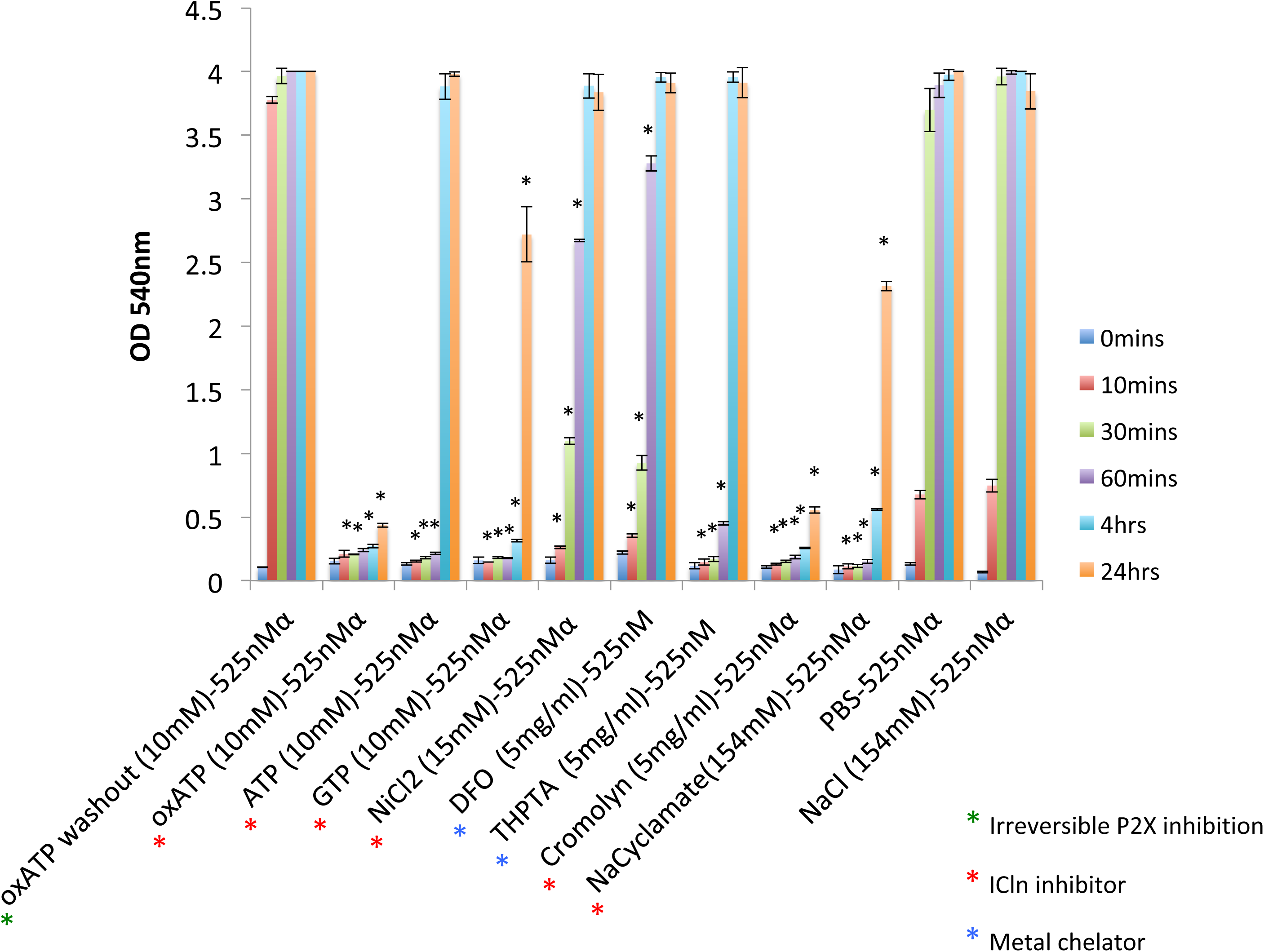
Inhibitors of the nucleotide-sensitive, heavy metal-binding ICln chloride channel block pore forming toxin-mediated hemolysis. **a)** Washed whole human blood was pre-incubated the chromone class of ICln inhibitors, cromolyn and nedocromil (5mg/mL) prior to ETX exposure (15nM at 37°C), and sampled over 24 hours. Data shown are from experiments performed in triplicate. Error bars represent standard deviations, and asterisks indicate that results are statistically significant compared with PBS vehicle control (green); Student’s *t*-test, \**P* < 0.0004. **b)** Washed whole human blood was pre-incubated with the ICln inhibitor, cyclamate anion (154mM) or acetate as a control organic anion (154mM) prior to ETX exposure (15nM at 37°C), and sampled over 24 hours. Data shown are from experiments performed in triplicate. Error bars represent standard deviations, and asterisks indicate that results are statistically significant compared with the NaCl control (green); Student’s *t*-test, \**P* < 0.0001. **c)** Washed whole human blood was pre-incubated with a full panel of known ICln inhibitors, ATP, oxATP, cromolyn (5mg/mL each); sodium cyclamate (154mM); and NiCl_2_ (15mM) prior to ETX exposure (15nM at 37°C), and sampled over 24 hours. Data shown are from experiments performed in triplicate. Error bars represent standard deviations, and asterisks indicate that results are statistically significant compared with the NaCl control (blue); Student’s *t*-test, \**P* < 0.0001. **d)** Human RBC membranes were analyzed for differential ICln expression in the setting of increasing ETX concentrations by Western blot analysis. Arrowheads signify individual components of the ICln monomer (closed) and its doublet (open). Data shown are from a single experiment and are representative of 3 independent experiments, using 3 distinct blood donors. **e)** Washed whole human blood was exposed to P2 receptor inhibition (green asterisks *), a full panel of ICln inhibitors (red asterisks *) and redox-active metal chelators (blue asterisks *) prior to being exposed to a hemolytic dose of *S. aureus* alpha-toxin (525nMα at 37°C) and sampled over 24 hours. Data shown are from experiments performed in triplicate. Error bars represent standard deviations, and black asterisks indicate that results are statistically significant compared with the NaCl as a control for NiCl_2_ and NaCyclamate, and PBS as a vehicle control for all nucleotides and metal chelators; Student’s *t*-test, \**P* < 0.0007.

Because other pore-forming toxins that form similarly sized pores, e.g., *Staphylococcus aureus* alpha toxin, have been reported to cause hemolysis via P2 receptor amplification [32], we explored the possibility that ICln may also be a target for other toxins. Indeed, when we pre-incubated human blood with a large panel of ICln inhibitors and heavy metal chelators, we found that *S. aureus* alpha toxin also employed the ICln channel, similar to what we had observed for ETX (Fig 4e).

### T cell lysosomal exocytosis contributes to ETX-mediated hemolysis

Because human T-lymphocytes also express MAL [33, 34], we also wanted to explore the possibility that T cells might contribute to ETX-mediated hemolysis. To assess possible leukocyte involvement, we compared the rate of ETX-mediated hemolysis in PBS-washed whole blood to that of PBS-washed RBCs that were leukocyte depleted (Fig 5a). The slowed rate of ETX-mediated hemolysis in the case of leukocyte depletion prompted to us to confirm that MAL-expressing CD4+ T cells do indeed bind ETX (Fig 5b). In agreement with the literature, ETX binding suggests that human T cells express MAL, while non-malignant B cells do not [35].

**Fig 5.**
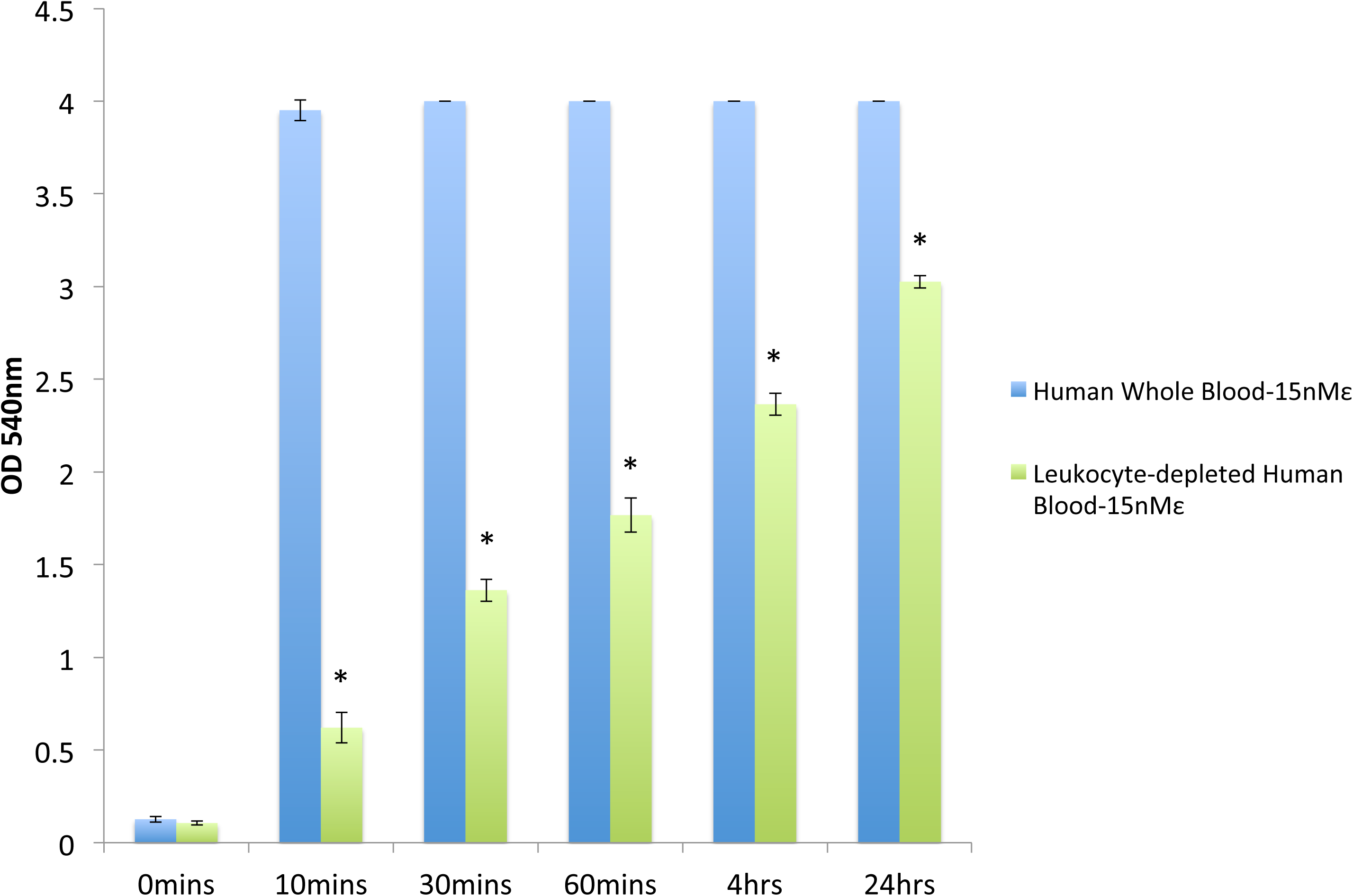
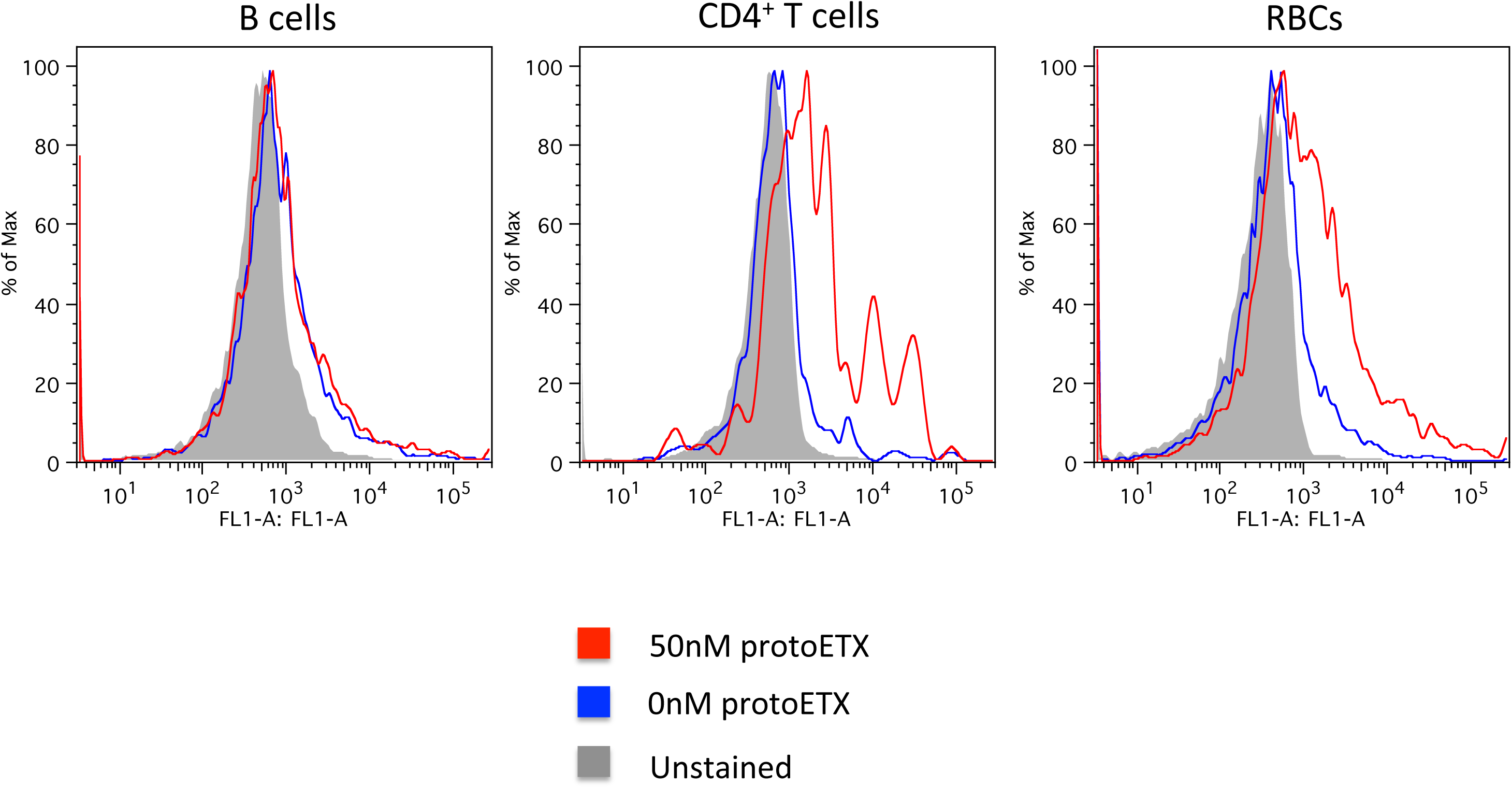
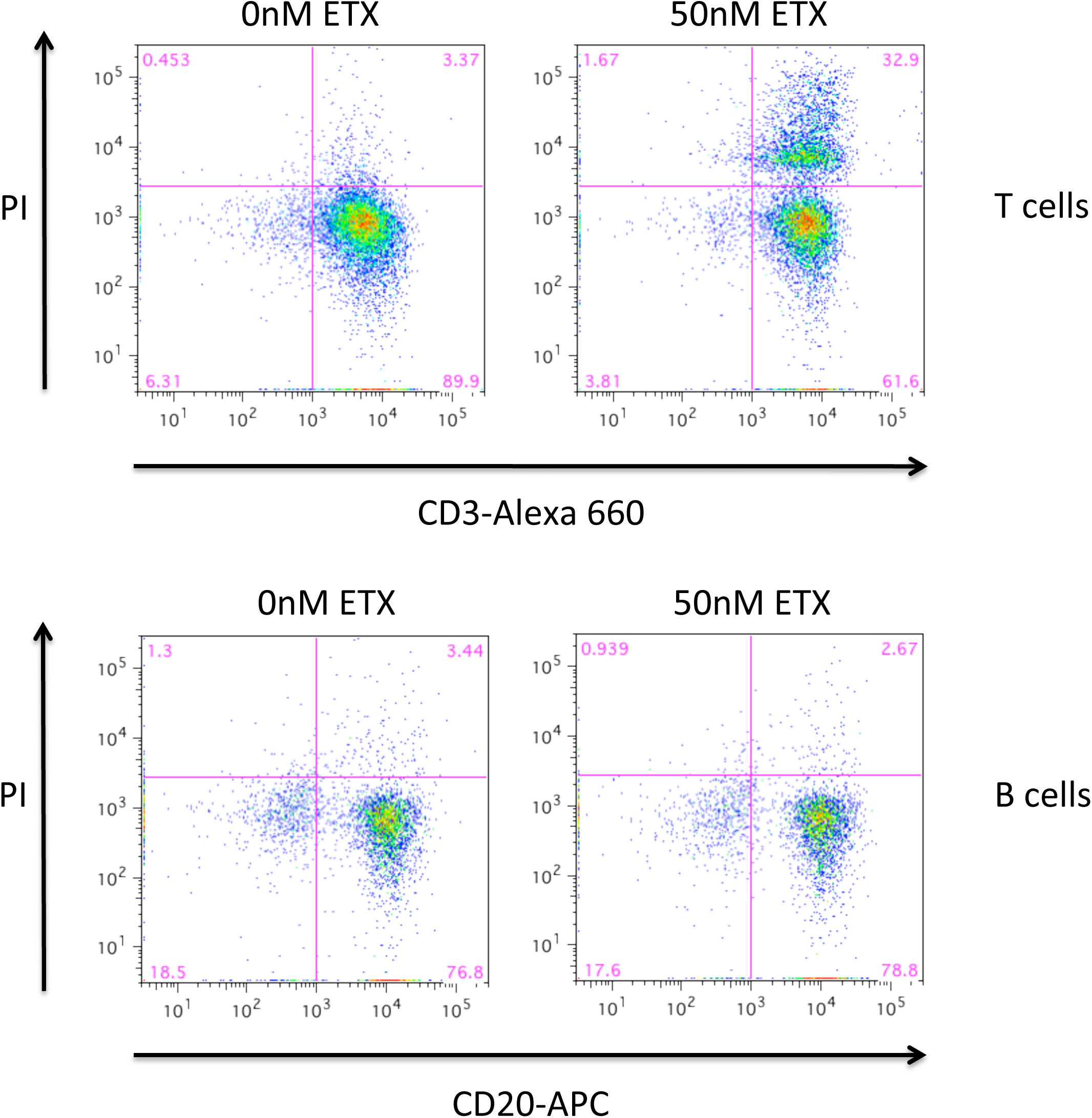
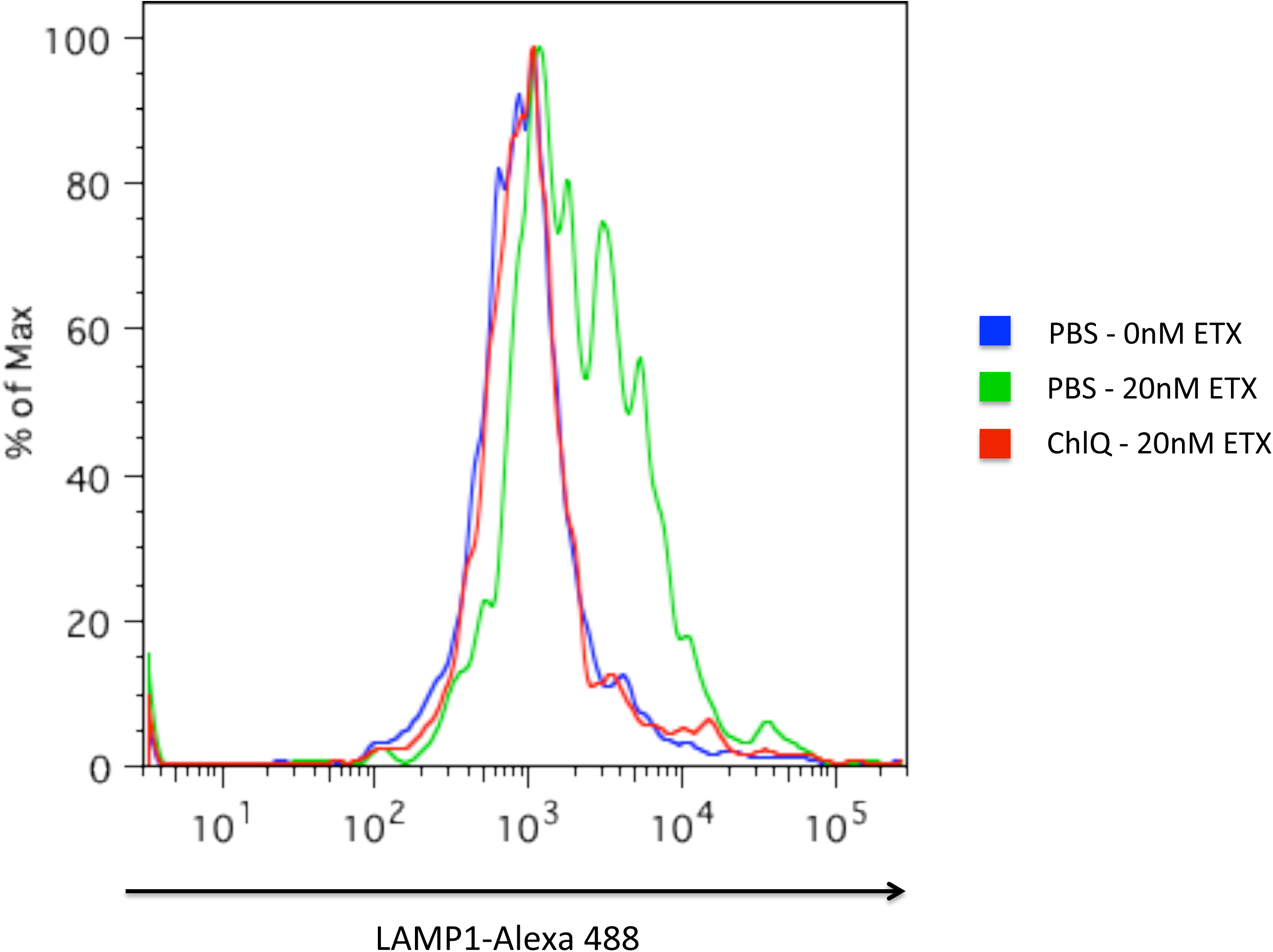
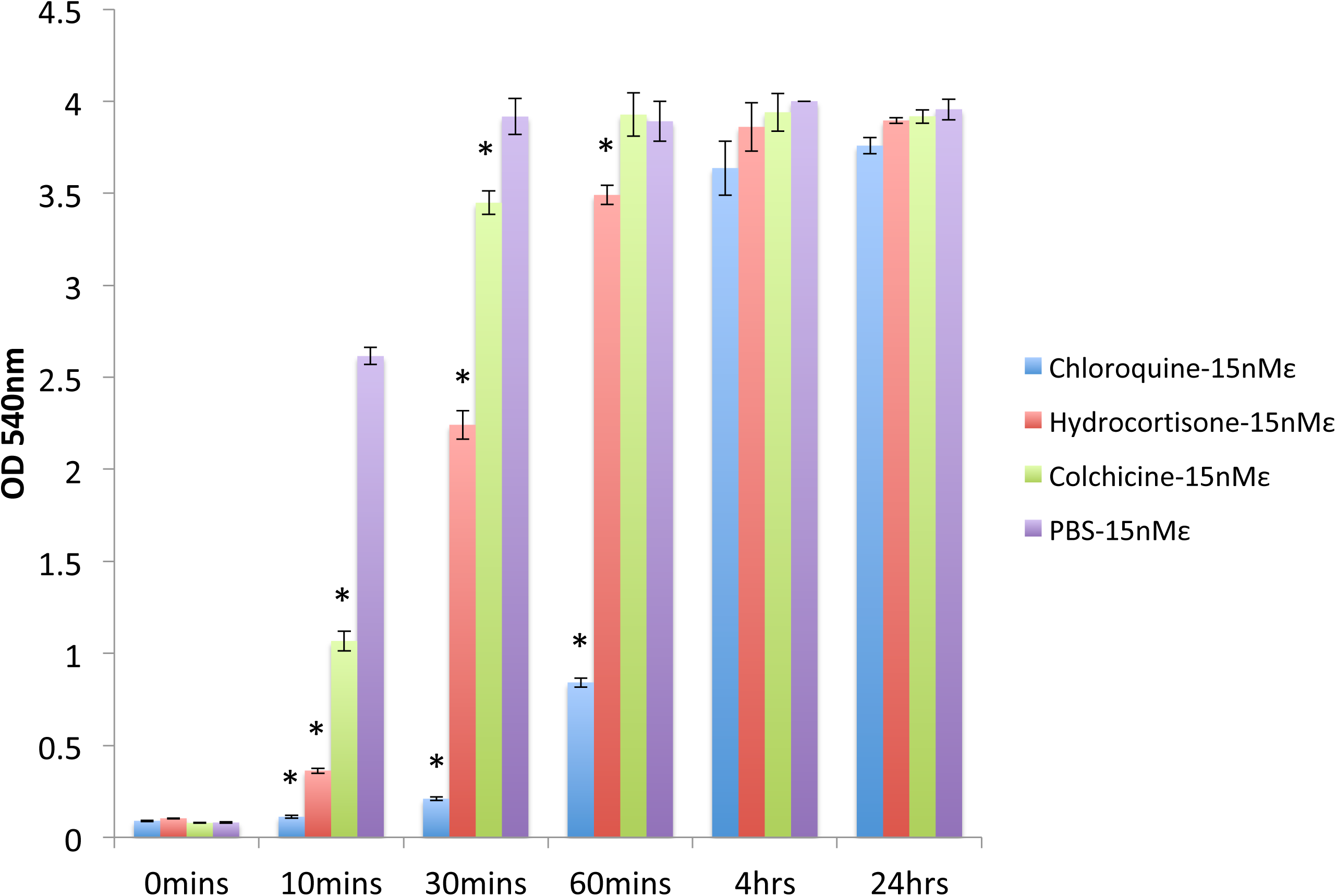
Leukocytes contribute to hemolysis via ETX-triggered T cell lysosomal exocytosis. **a)** Leukocyte-depleted human blood was washed and suspended in PBS, incubated with 15nM ETX at 37°C and sampled over 24 hours. Hemolysis was compared to that of washed whole human blood incubated under the same experimental conditions. Data shown are from experiments performed in triplicate. Error bars represent standard deviations, and asterisks indicate that results are statistically significant compared with human blood (blue); Student’s *t*-test, \**P* < 0.0001. **b)** Isolated B cells, T cells and human RBCs were incubated with non-toxic protoETX (50nM) for 2 hours at 4°C. Toxin binding was detected via flow cytometry using custom anti-ETX antibody, JL001.2. Data shown are from a single experiment and are representative of 3 independent experiments, using 3 distinct blood donors. **c)** Isolated T and B cells were exposed activated ETX (50nM) for 1 hour at 37°C and assessed for membrane integrity by propidium iodide (PI) uptake. Data shown are from a single experiment and are representative of 3 independent experiments, using 3 distinct blood donors. **d)** Isolated T cells were exposed to activated ETX (20nM) for 1 hour at 37°C with or without the lysosomotropic agent, chloroquine (ChlQ). Data shown are from a single experiment and are representative of 3 independent experiments, using 3 distinct blood donors. **e)** Washed whole human blood was suspended in PBS and incubated with different inhibitors of lysosomal exocytosis, chloroquine, hydrocortisone hemisuccinate and colchicine, 4 hours prior to ETX administration. All samples were then treated with ETX (15nM), incubated at 37°C, and sampled over 24 hours. Data shown are means and SD from a single experiment and are representative of 3 independent triplicate experiments. Asterisks indicate that results are statistically significant compared with PBS vehicle control (purple); Student’s *t*-test, \**P* < 0.005.

Blanch et al. have recently determined that ETX forms pores in the membranes of a human T-lymphocyte cell line [36]. Moreover, they demonstrated that T-lymphocyte cell lines are sensitive to ETX toxicity, while B-lymphocyte cell lines are not. We wished to confirm that primary T cells are indeed susceptible to ETX. Therefore, we isolated CD3+ T cells and exposed them to ETX (50nM) for 1 hour at 37°C and assessed cell death by propidium iodide (PI) uptake. Isolated CD20+ B-lymphocytes were used as a control lymphocyte population. Results revealed that a subset of human T cells is sensitive to ETX-mediated damage, while human B cells are completely refractory (Fig 5c).

One way that nucleated cells can defend against pore-forming toxins is via lysosomal exocytosis [37, 38]. However, this pathway has been described for toxins that form relatively large pores (> 2nm in diameter) in the plasma membrane. For example, streptolysin O allows the influx of Ca^2+^, which then triggers the lysosomal exocytosis pathway and lysosomal hydrolase-mediated membrane internalization and repair [37, 39]. Toxins similar to ETX that form much smaller pores, such as *Staphylococcus aureus* alpha toxin, have not previously been reported to trigger lysosomal exocytosis. Nucleated cells exposed to small pore formers are thought to first internalize these toxins and then expel them in a membrane-bound exosome-like form [40]. However, the exocytosis machinery has not been clearly identified. Along these lines, we wished to determine if the lysosomal exocytosis pathway might be involved in how nucleated cells, e.g., human T cells, process small pore-forming toxins such as ETX. Intriguingly, similar to *Staphylococcus aureus* alpha toxin, ETX has been reported to undergo internalization soon after binding the plasma membrane of nucleated cells [41].

Administration of ETX (20nM) to human T cells caused significant surface expression of lysosome associated membrane protein-1 (LAMP1), a marker for the limiting membrane of the lysosome, and thus a marker for lysosomal exocytosis [39]. To determine if normal lysosomal function is required for ETX-mediated lysosomal exocytosis, we disrupted lysosomal acidification and subsequent function, by exposing ETX-treated T cells to chloroquine, a lysosomotropic agent that strongly suppressed ETX-mediated lysosomal exocytosis (Fig 5d). Chloroquine preferentially accumulates in acidic compartments of the cell and prevents acidification because of its protonated basic amine groups [42, 43].

Remarkably, lysosomal exocytosis has previously been proposed as a hemolytic mechanism in the case of anti-Rh sensitized RBCs and engaging monocytes [44]. In this previous study, the drugs colchicine (a microtubule depolymerizing agent), and hydrocortisone hemisuccinate (a corticosteroid), suppressed monocyte-mediated hemolysis. The authors suggested that the anti-hemolytic effect of these drugs was the result of suppressing monocyte lysosomal exocytosis [45]. Similarly, we wished to determine if blockade of lysosomal exocytosis by chloroquine or other inhibitory compounds, colchicine and hydrocortisone hemisuccinate, was sufficient to inhibit ETX-mediated hemolysis. Indeed, we found that each of these compounds inhibited ETX-mediated hemolysis, chloroquine > hydrocortisone > colchicine (Fig 5e).

## DISCUSSION

We have confirmed the findings of Gao et al. that human blood exposure to *C. perfringens* epsilon toxin results in hemolysis. However, our data extend these findings and suggest that metal-catalyzed oxidation of the nucleotide-sensitive, volume-regulating ICln channel is likely responsible for amplifying the hemolytic process, rather than P2 receptor activation. Moreover, we find that ICln may be involved in hemolytic amplification for pore-forming toxins beyond ETX, e.g., *S. aureus* alpha toxin and perhaps others. An important experimental distinction between this study and that of Gao et al. is the substantial difference in the concentration of ETX used. Our hemolytic assays were conducted at 15nM ETX, while Gao et al. used ≥ 100nM ETX [20].

We have also shown that human RBCs express a minor MAL isoform (likely MAL isoform C), which would explain why human RBCs are uniquely susceptible to ETX-mediated damage, as compared to RBCs from other species that do not express MAL. Similarly, ETX also damages MAL-expressing human T cells, causing them to undergo lysosomal exocytosis. Our data also show that lysosomal exocytosis contributes to ETX-mediated hemolysis, as evidenced by hemolytic blockade by lysosomal inhibitors such as chloroquine. Considering chloroquine’s ability to increase pH in a general fashion, we suspect that it may have a dual effect on inhibiting ETX-mediated hemolysis, as ICln’s ion conductance is pH sensitive [27]. Along these lines, we have observed that conducting hemolysis assays in bicarbonate buffer also results in slowed ETX-mediated hemolysis (data not shown). Therefore, the inhibitory action of hydrocortisone and colchicine might more accurately reflect the true contribution of T cell lysosomal exocytosis.

To better understand how lysosomal exocytosis might influence ETX-mediated hemolysis, we reviewed the literature and identified that lysosomes serve as storage compartments for redox-active heavy metals such as Cu^+^ and Fe^3+^[46]. For example, the copper transporter ATP7B actively transports Cu^+^ into the late endosome, which later fuses with the lysosome [47]. Furthermore, an excess of extracellular copper stimulates lysosomal copper uptake, and the cell can go on to release accumulated copper into the extracellular space via expulsion through the lysosomal exocytosis pathway [48]. For these reasons, we favor the idea that ETX triggers the release of previously stored redox-active heavy metals from the lysosomal compartment of MAL-expressing T cells, which are predominantly of the CD4+ lineage [35]. To help visualize this process, we have composed a diagram to illustrate the proposed mechanism in a step-wise fashion (Fig 6), while figure 7 aims to illustrate a more comprehensive schema for ETX-mediated hemolysis.

**Fig 6.**
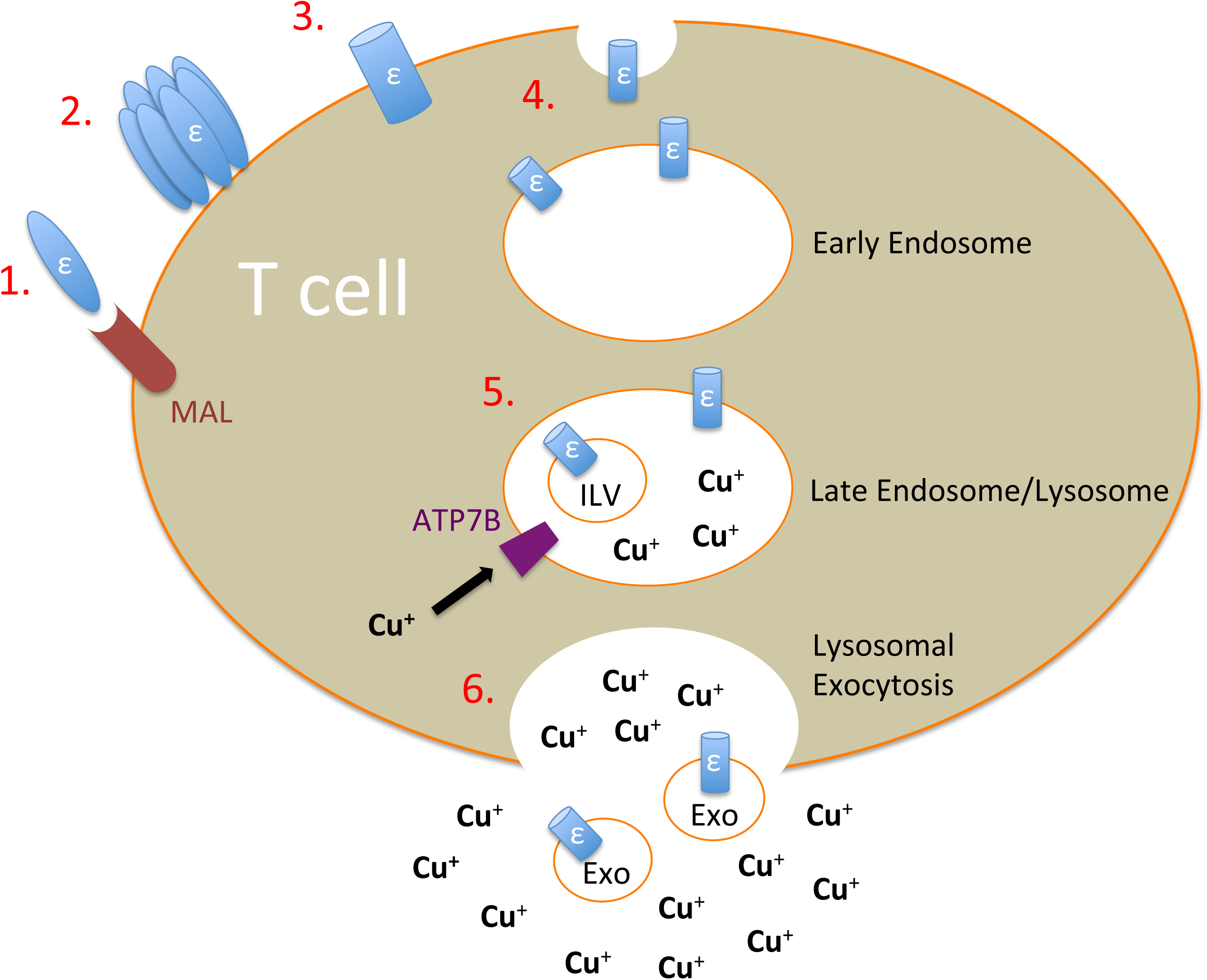
A schematic illustration of ETX-mediated lysosomal exocytosis. **1)** Epsilon toxin (ε) binds to the T cell plasma membrane via its cellular receptor (possibly Myelin And Lymphocyte protein, MAL). **2)** Seven ETX subunits form a pre-pore complex at the cell surface. **3)** The heptameric pre-pore matures into a transmembrane pore (cylinder). **4)** Pore formation triggers membrane invagination and endosome formation. **5)** As the late endosome matures, ETX-containing intraluminal vesicles (ILV) begin to form. Concurrently, cytosolic Cu^+^ ions are being transported into the late endosome via the ATP7B copper transporter. **6)** Upon becoming a mature lysosome, fusion occurs between the lysosomal membrane and the plasma membrane, resulting in lysosomal exocytosis. Lysosomal exocytosis releases exosome-like, ETX-containing microparticles (Exo), similar to what has previously been described for toxins that form small pores (< 2nm in diameter) such as *S. aureus* alpha toxin [40], into the extracellular space. In addition to releasing ETX-containing microparticles, redox-active heavy metals stored in the lysosome such as Cu^+^ are also released.

**Fig 7.**
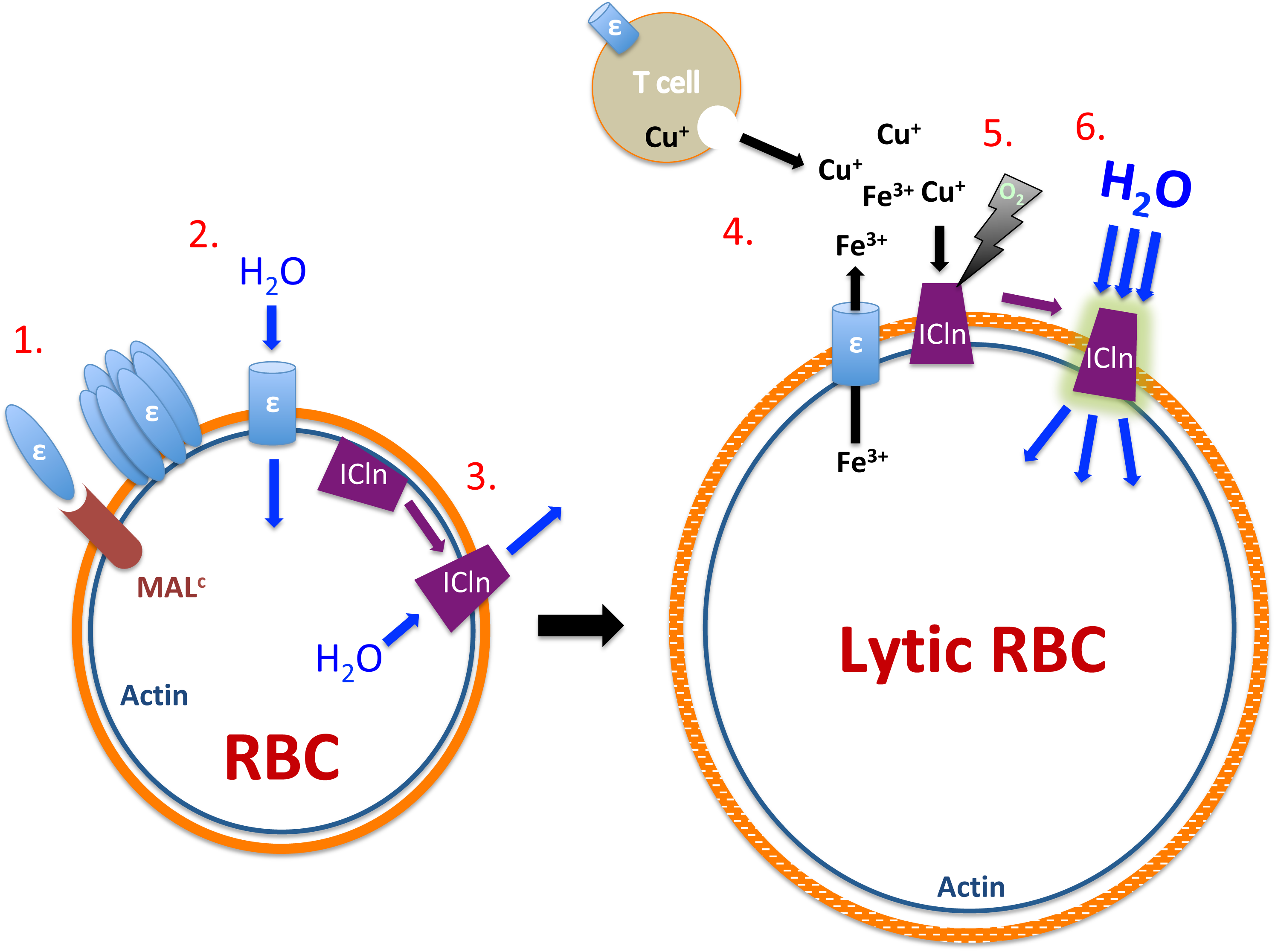
A schematic illustration of ETX-mediated RBC lysis. **1)** Epsilon toxin (ε) binds to the human RBC membrane via a cellular receptor (possibly MAL isoform C). **2)** Oligomerized toxin forms a pore and inserts into the RBC membrane, allowing unregulated influx of ions and water. **3)** Cellular swelling causes rapid insertion of the ICln chloride channel as a countermeasure to resist osmolysis, resulting in the efflux of ions and water. **4)** The ETX pore facilitates the release of intracellular redox-active heavy metals from both the T cell (Cu^+^) and RBC cytosol (Fe^3+^). **5)** Extracellular heavy metals bind to and damage the ICln pore in an oxygen-dependent fashion. **6)** Metal-catalyzed oxidation of the ICln pore results in a deregulated channel that allows ions and water to flow down their diffusion gradient resulting in eventual osmolysis.

The notion that ICln must first insert into the RBC membrane for metal-catalyzed oxidation to ensue might lead to the prediction that supernatant harvested from ETX-exposed T cells would not be sufficient to trigger ETX-mediated hemolysis, and indeed, we find this to be the case (Supp Fig 4). However, an important caveat to this experiment is that Cu^+^ ions are unstable in aqueous solution due to rapid oxidation (oxygen-dependent) and rapid disproportionation (oxygen-independent), and thus must be chelated in the cytosol by chaperones such as glutathione that are largely absent in the extracellular space [49]. This lack of extracellular stability could lead to a proximity requirement for toxin-exposed T cells to have a hemolytic effect on neighboring RBCs. To address this caveat, we transplanted isolated human leukocytes into blood harvested from non-human species, i.e., cow and rhesus macaque; RBCs that have been shown to either lack MAL (cow) or that fail to bind ETX (macaque). ETX-sensitive human leukocytes were unable to trigger hemolysis in non-human RBCs, even when exposed to 150nM ETX (Supp Fig 5).

In its entirety, this study may help to shed light on how RBCs incur damage during an acute MS relapse because ETX, a blood-borne neurotoxin, has previously been indicated as a potential MS trigger due to its remarkable specificity for BBB vasculature and for CNS myelin [9, 10]; the two tissues damaged during each MS attack. The finding that ETX also causes RBC abnormalities, reminiscent of what has been observed during active MS, may further support the ETX-MS hypothesis. Of note, even though we assessed ETX-mediated hemolysis in > 70 study participants, we did not observe any ETX-resistant phenotypes, suggesting that all blood types tested were ETX-susceptible and express MAL. To our knowledge, MAL expression by circulatory cells has not been reported in any non-human species to date, thus blood cell sensitivity to ETX may be an exclusively human trait.

Should ETX be conclusively shown to trigger nascent MS lesion formation, ICln insertion into the RBC plasma membrane and surface LAMP1 expression on CD4+ T cells may lay the groundwork for novel biomarker development. To date, there are no biomarkers that can predict MS disease activity. The identification of such biomarkers may be useful in predicting the onset of neurological symptoms and/or identifying occult disease activity, as is the case for “silent lesions,” which are detectable on MRI but yield no observable symptoms [50].

In addition to surface LAMP1 expression, lysosomal exocytosis also causes the release of lysosomal hydrolases into the extracellular milieu, and the enzymatic activity of acid hydrolases such as β-hexosaminidase are commonly used as markers for this cellular process [48]. These enzymes may also be candidate biomarkers for ETX blood exposure and early disease activity in multiple sclerosis.

Finally, identifying the mechanism by which ETX causes RBC lysis may allow for novel clinical interventions. For example, metal chelation therapy might inhibit hemolysis, thus preventing vascular iron deposition and neuronal toxicity/axonal loss. Although hemolysis and free hemoglobin have recently been shown to correlate with the transition from RRMS to SPMS [8], the root cause of this hemolysis has not yet been identified. *C. perfringens* epsilon toxin may adequately explain this phenomenon, in addition to how nascent MS lesions form in the absence of an inflammatory infiltrate [9, 51].

## MATERIALS AND METHODS

### Ethics Statement

Research protocol RRU-0952 for the collection of samples from individuals with MS and healthy controls was reviewed and approved by the Rockefeller University institutional review board. All participants in the study gave written informed consent.

### Epsilon Toxin

His-tagged protoxin was procured from BEI Resources, activated by adding an equal volume of 0.25% trypsin (ThermoFisher) and incubating for 1 hour at 37°C. Trypsin was then inactivated by the addition of 1:1 volume Defined Trypsin Inhibitor (ThermoFisher).

### Blood Sample Collection and Manipulation

Human blood from healthy adult donors was obtained at the Rockefeller University hospital using heparinized tubes. Cow, goat, sheep, rat, guinea pig and rhesus macaque blood and enriched human, cow and rat RBCs were all purchased from Innovative Research, Inc. All whole blood and enriched RBC samples were centrifuged for 5 mins at 600g, the supernatant was aspirated, and the cell pellet was washed with PBS (20x the original volume) prior to re-suspension in PBS so as to match the original starting blood volume. For experiments using transition metals nickel and manganese, human blood was washed as previously described. However, normal saline was used instead of PBS to avoid the formation of insoluble Ni_3_(PO_4_)_2_ and Mn_3_(PO_4_)_2_ salts.

For the transfer of human leukocytes to the blood of non-human mammals, human blood was centrifuged at 600 x g for 5 mins and the plasma layer removed. Pelleted cells were suspended in 50 volumes of phosphate buffered saline (PBS), centrifuged at 1000 x g, and suspended in RBC lysis buffer (155 mM NH_4_Cl, 12 mM NaHCO_3,_ 0.1 mM EDTA). RBCs were allowed to lyse for 5 mins, after which, leukocytes were centrifuged at 600 x g for 5 mins. Pelleted leukocytes were washed 3 times in PBS and transferred to washed whole cow or whole macaque blood of a similar original volume.

### Hemolysis Quantitation

After ETX incubation, cells were pelleted at 600g for 5mins, supernatant was harvested and the degree of hemolysis was determined by light absorbance (OD 540nm) using a SpectraMax M5 Multi-Mode Microplate Reader (Molecular Devices).

### Lymphocyte Isolation

Harvested human blood was incubated with RosetteSep Human T cell enrichment cocktail (STEMCELL Technologies) or RosetteSep Human B cell enrichment cocktail (STEMCELL Technologies) and then isolated with Ficoll-Paque Plus (GE Healthcare), as per the manufacturer’s instructions.

### Flow Cytometry Staining and Analysis

For protoETX binding, isolated cells were washed 3 times with PBS 1% BSA buffer, and incubated for 2 hours at 4°C with 5% donkey serum (PBS 1% BSA) containing either 50nM or 0nM protoETX. Cells were washed 3 times with chilled PBS 1% BSA buffer, and then stained for 1 hour at 4°C with primary antibodies: rabbit anti-ETX (JL001.2, 1:1000), mouse anti-CD3 Alexa 660 (eBioscience) 1:200 or mouse anti-CD20 APC (eBioscience) 1:200. After 3 washes with chilled PBS 1% BSA, cells were then incubated with Alexa 488-conjugated donkey anti-rabbit (1:1000, Jackson ImmunoResearch) so as to detect bound JL001.2 antibody, and then thoroughly washed prior to Flow Cytometry analysis. For cytotoxicity assays, isolated lymphocytes were stained with mouse anti-CD3 Alexa 660 (1:200) or mouse anti-CD20 APC (1:200), and propidium iodide (ThermoFisher) 1:500. For lysosomal exocytosis assays, isolated T cells were stained with mouse anti-LAMP1 Alexa 488 (ThermoFisher) 1:200. All cells were analyzed by Flow Cytometry with BD AccuriC6 at our core facility.

### Western blot Analysis of RBC membrane proteins

100µL of purified RBCs were lysed in 900µL ammonium chloride RBC lysis buffer for 10 mins at 37°C. The cell lysate was centrifuged at > 16,000g for 5 mins, and the pellet was washed 3 times in PBS and re-suspended in 50µL PBS. An equal volume of 2X Laemmli sample buffer (Bio-Rad) was added to each sample and the dissolved membranes were stored at −20°C. Thawed samples were diluted 2.5 fold in 1X Laemmli sample buffer prior to SDS PAGE. Proteins were transferred to an Immobilon-P membrane (MilliporeSigma) and probed with either mouse-anti MAL 1:1000 (clone E-1, Santa Cruz Biotechnology, Inc.) or rabbit anti-ICln 1:1000 (PA5-13450, ThermoFisher). HRP-conjugated donkey anti-mouse IgG and donkey anti-rabbit IgG secondaries (Jackson Immunoresearch) were used with the corresponding primary antibody to visualize MAL expression and ICln expression respectively (1:100,000). SuperSignal West Femto Maximum Sensitivity Substrate (ThermoFisher) was used to visualize bound antibody.

### Anaerobiosis

Anaerobic experiments were performed in an anaerobic hood (BACTRON Anaerobic Chamber).

### Drugs and Compounds

All drugs and compounds used in this study were purchased from Sigma Aldrich.

### Statistical Analysis

Results are representative of data obtained from repeated independent experiments. Each value represents the mean ± *SD* for three replicates. Statistical analysis was performed using the two-tailed Student *t*-test (GraphPad Software, San Diego, CA, USA).

## ACKNOWLEDGEMENTS

We would like to thank Dr. Timothy Vartanian and Dr. Jennifer Linden for providing the anti-ETX detection antibody, JL001.2, and the anti-ETX neutralizing antibody, JL008.

---

**Supplemental Fig 1. SDS PAGE analysis of His-tagged protoETX**

His-tagged prototoxin (BEI Resources) was diluted in PBS and combined with 2X laemmli sample buffer (1:1 volume), heated at 90°C for 6 minutes and subjected to electrophoresis on a 4-12% Bis-Tris gel. Proteins were stained using a colloidal blue protein staining kit.

**Supplemental Fig 2. ProtoETX requires trypsin activation to gain hemolytic activity.**

Washed whole human blood was exposed to trypsin activated ETX (15nM) or non-toxic protoETX (15nM), incubated at 37°C, and sampled over 24 hours. Data shown are means and SD from a single experiment and are representative of 3 independent triplicate experiments. Asterisks indicate that results for protoETX (blue) are significant different when compared to active ETX (red); Student’s *t*-test, \**P* < 0.0001.

**Supplemental Fig 3. Neutralizing anti-ETX monoclonal JL008 inhibits ETX-mediated hemolysis in a dose-dependent manner. a)** Washed whole human blood was pre-incubated with a neutralizing anti-ETX rabbit monoclonal (JL008) at varying concentrations (10^-2^, 10^-3^ and 10^-4^) and compared to a rabbit isotype control antibody (Rb IgG), and PBS vehicle control. All samples were treated with ETX (15nM), incubated at 37°C, and sampled over 24 hours. Data shown are means and SD from a single experiment and are representative of 3 independent triplicate experiments. Asterisks indicate that results are statistically significant compared with PBS vehicle control (light blue); Student’s *t*-test, \**P* < 0.0007.

**Supplemental Fig 4. Supernatant from ETX-exposed human T cells is not sufficient to trigger hemolysis.** Human T cell isolates were treated with ETX (30nM) at 37°C for 1 hour in PBS buffer (1% BSA). Supernatant was harvested and unbound ETX was neutralized with anti-ETX rabbit monoclonal, JL008 (10^-2^). Alternatively, the supernatant was incubated with a control rabbit isotype control antibody, Rb IgG (10^-2^). Supernatants were added to washed whole human blood and incubated at 37°C, and sampled over 24 hours. Data shown are means and SD from a single experiment and are representative of 3 independent triplicate experiments. Asterisks indicate that results are statistically significant compared with the positive control, 30nM ETX (purple); Student’s *t*-test, \**P* < 0.0001.

**Supplemental Fig 5. Transfer of human leukocytes into non-human, ETX-resistant blood, fails to confer ETX hemolytic sensitivity.** Human leukocytes were isolated and transferred into washed whole cow or whole rhesus macaque blood, incubated with 150nM ETX at 37°C, and sampled over 24 hours. Data shown are means and SD from a single experiment and are representative of 3 independent triplicate experiments. Asterisks indicate that results are statistically significant compared with ETX (150nM) treated, washed whole human blood (blue); Student’s *t*-test, \**P* < 0.0001.

